# Interaction with tumor cell spheroids induces suppression in primary human cytotoxic T cells

**DOI:** 10.1101/2025.03.18.643973

**Authors:** Amal Alsubaiti, Hanin Alamir, Lan Huynh, Tressan Grant, Abdullah Aljohani, Po Han Chou, Yiwei Shi, Maryam Alismail, Lydia R. Mason, Andrew Herman, John S. Bridgeman, Christopher J. Holland, Christoph Wülfing

## Abstract

Cytotoxic T lymphocytes (CTL) are key effectors in the anti-tumor immune response. However, their function is commonly suppressed in tumors in the form of exhausted CTL. Understanding molecular mechanisms of suppression and of therapeutics to overcome them is of substantial basic and translational importance yet hindered by limited access to large numbers of exhausted CTL in vitro. Here we use three-dimensional tissue culture to generate primary human CTL with suppressed function. Using a 21-antibody flow cytometry panel and determination of calcium signaling and cell couple maintenance, we show that these cells closely resemble exhausted CTL from tumors. For better understanding of in vitro human primary CTL as key tools in therapeutic development, before and after induction of suppression, we have determined the dependence of CTL function on technicalities of in vitro CTL generation, antigen dose and affinity across two T cell receptors and multiple tumor cell lines. We have investigated morphology and subcellular F-actin distributions of CTL as a key regulators of effector function. Primary human CTL formed cell couples with tumor target cells even in the absence of antigen. Yet, gradual stabilization of such cell couples was associated with increasing CTL effector function. Induction of suppression substantially destabilized CTL tumor cell couples. This comprehensive characterization of the phenotype of in vitro primary human CTL, including a suppressed state, should facilitate their wider use in basic and translational research.

## Introduction

Cancer is frequently associated with an anti-tumor immune response that involves most immune cell types. CD8^+^ cytotoxic T lymphocytes (CTL) have the inherent ability to kill tumor target cells, yet this ability is commonly suppressed in the tumor microenvironment. Understanding mechanisms of CTL suppression and how therapeutics can overcome them is of wide interest. Such research greatly benefits from efficient, scalable access to suppressed CTL. Omics-based analyses of patient samples have established that CTL suppression consists of multiple progressive CTL states from precursor exhausted to terminally exhausted CTL (Beltra et al. 2020, Giles et al. 2023, Hudson et al. 2023, Mellman et al. 2023). Any model of suppressed CTL needs to map onto this in vivo progression.

Mouse models provide access to suppressed CTL and have greatly contributed to the understanding of anti-tumor immune responses (Guerin et al. 2020). However, tumor development is commonly accelerated in mice, keeping mice under specific pathogen-free conditions as universally done alters key aspects of immune function (Rosshart et al. 2019) and using mice in therapeutic development requires the generation of matched murine and human compounds. Humanized mice are available yet require substantial genetic engineering (Allen et al. 2019). Therefore, effective access to human suppressed CTL is of substantial utility. Patient-derived organoids that retain some immune function or can shape CTL function are now available (Dijkstra et al. 2018, Jenkins et al. 2018, Yuki et al. 2020, Polak et al. 2024). They can provide an in vitro model of the interaction of the entire immune system with a tumor, yet application at scale is difficult. Organ-on-chip models offer the potential of scale (Pavesi et al. 2017, Park et al. 2019, Kerns et al. 2021, Maulana et al. 2021), yet their validation against in vivo biology can be challenging.

Here we present a scalable experimental approach to generate primary human suppressed CTL that closely resemble tumor-infiltrating CTL. In previous murine work we have shown that the phenotype of CTL that have interacted with tumor cells presenting antigen and grown as three-dimensional spheroids closely resembles the phenotype of the same CTL interacting with the same tumor cells grown subcutaneously in vivo (Ambler et al. 2020). Here we have transferred this strategy to primary human cells. Human CTL were grown from peripheral blood mononuclear cells (PBMC) and lentivirally transduced to express the 1G4 or MEL5 T cell receptor (TCR) recognizing peptides derived from the tumor associated antigen New York esophageal squamous cell carcinoma 1 (NY-ESO-1) or melanoma antigen A(Melan-A)/melanoma antigen recognized by T cells 1 (MART-1), respectively (Chen et al. 2005, Madura et al. 2019). We have characterized agonist peptide-dependent killing and IFNγ secretion as a function of technicalities of CTL generation and across peptide concentrations and affinities in response to Mel624 or A375 melanoma and NCI-H1755 non-small cell lung carcinoma cells. For effective killing CTL need to undergo a series of subcellular polarization steps that cumulate in the release of lytic granules (Ritter et al. 2015, Ambler et al. 2020). We have characterized the execution of such polarization as a function of stimulus strength and shown that gradual stabilization of CTL tumor cell couples was associated with increasing CTL effector function.

Using a 21-antibody flow cytometry panel, we have shown that human CTL grown in vitro from PBMC have a partially suppressed phenotype. Interaction of these human CTL with tumor cell spheroids induced further suppression, as evident in substantially reduced effector function, impaired subcellular polarization, loss of calcium signaling and a flow cytometry phenotype that closely resembles exhausted CTL from tumors. This work thus provides access to primary human suppressed CTL of an extensively characterized phenotype. These cells allow for efficient and physiological investigation of mechanisms of CTL suppression and of mechanisms of action of therapeutics, as already initiated (Alamir et al. 2023).

## Results

### Primary human CTL expressing a transgenic TCR kill tumor target cells

The generation of suppressed primary human CTL required two steps, the generation of primary human CTL expressing a transgenic TCR and the subsequent interaction of such cells with tumor cell spheroids. To generate primary human CTL expressing a transgenic TCR, we isolated CD8^+^ T cells from buffy coats and activated these cells with beads coated with antibodies against CD3ε and CD28. We transduced these cell cultures with a human immunodeficiency virus (HIV)-derived, vesicular stomatitis virus (VSV) G protein-pseudotyped lentivirus driving the expression of the alpha and beta chains of a transgenic TCR and GFP as a transduction marker (Fig. 1A). Transcription was driven by a spleen focus-forming virus (SFFV) promoter (Morsut et al. 2016). The three proteins were linked by 2A ribosome skipping signals (Szymczak et al. 2004). Furin cleavage sites allowed for the cleavage of the 2A peptides after translation. We used two TCRs. The 1G4 TCR recognizes peptide 157-165 of the tumor-associated antigen NY-ESO-1 presented by HLA-A*0201 (Chen et al. 2005). The MEL5 TCR recognizes peptide 26-35 of the tumor-associated MART-1 also presented by HLA-A*0201 (Madura et al. 2019). For both TCRs, pairing of the transgenic alpha and beta chains was promoted by the addition of a disulfide bridge between the constant domains (Cohen et al. 2007) as used consistently here. For the MEL5 TCR we separately compared promotion of pairing of the transgenic alpha and beta chains through a disulfide bridge to that through use of murine constant domains (Cohen et al. 2006) as further described below.

**Figure 1.**
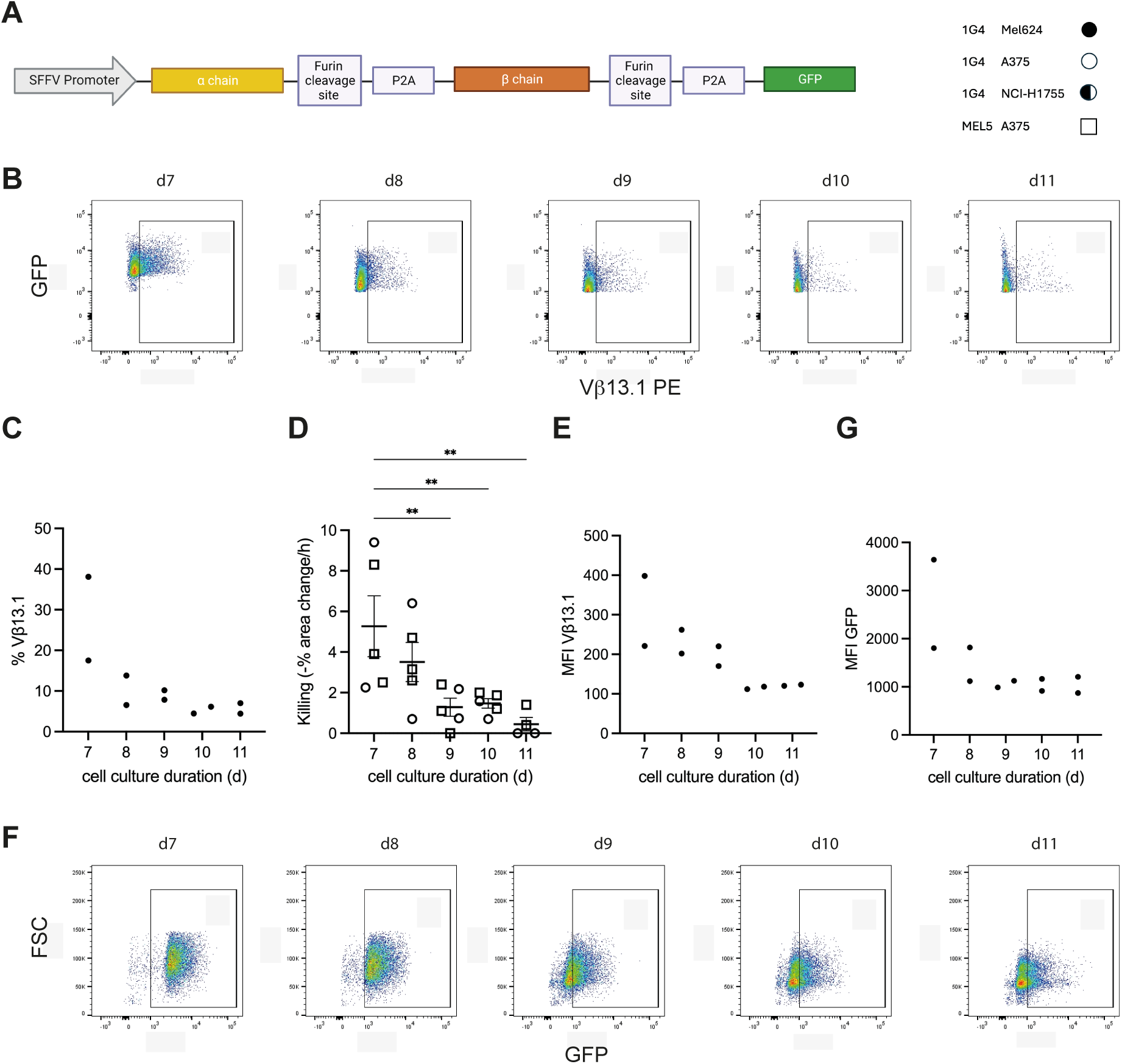
Transient expression of transgenic TCR in human primary CTL **A** The lentiviral cassette for the expression of the transgenic TCR and symbols used across all figures for the interaction of CTL transduced to express a transgenic TCR, 1G4 or MEL5, with indicated tumor target cell lines. **B** Representative staining data of human CTL transduced to express the 1G4 TCR with GFP as the transduction marker for the Vβ13.1 element of the 1G4 TCR after the indicated number of days of cell culture. Shown are only live cells that were sorted for high GFP expression, GFP^+++^. The gate to identify Vβ13.1-positive CTL is indicated. 1 of 2 independent experiments. **C** Percent CTL transduced to express the 1G4 TCR in the GFP^+++^ sort gate that are positive for Vβ13.1 after the indicated number of days of cell culture. 2 independent experiments. **D** Killing of A375 tumor target cells by CTL transduced to express the 1G4 or MEL5 TCR and sorted for high GFP expression after the indicated number of days of cell culture as mean ± SEM. 5 independent experiments. Statistical significance determined by paired One-way ANOVA. **E** Vβ13.1 MFI of the same cells as in C. **F** Representative GFP fluorescence data of human CTL transduced to express the 1G4 TCR with GFP as the transduction marker after the indicated number of days of cell culture. All live cells are shown. The gate to identify GFP-positive CTL is indicated. 1 of 2 independent experiments. **G** GFP MFI of the same cells as in C. 2 independent experiments. ** p<0.01.

To evaluate cytolytic function of the human primary CTL expressing a transgenic TCR, we used an imaging-based cytotoxicity assay (Ambler et al. 2020). We used three target cell lines expressing HLA-A*0201, the melanoma cell lines Mel624 and A375 and the non-small cell lung carcinoma cell line NCI-H1755. For the initial characterization of the CTL, the cell lines were incubated with a high concentration (2µg/ml) agonist peptide, NY-ESO-1157-165 and MART-126-35 with the A27L mutation (‘ELA’) that enhances MHC binding (Madura et al. 2019). Endogenous antigen presentation and peptide dose responses are characterized below. For use in the imaging-based killing assay, the cell lines were stably transfected to express the red fluorescent protein mCherry (Shaner et al. 2004). Lentivirally transduced CTL were cultured for 7-11 days in the presence of IL-2 plus beads coated with antibodies against CD3ε and CD28, as further explored below. Transgenic TCR expression was determined with an antibody against the Vβ element of the TCR, Vβ13.1 for the 1G4 TCR and Vβ20 for the MEL5 TCR. TCR expression was moderate, not exceeding 40% TCR Vβ-positive in CTL sorted for GFP expression of at least 10-fold above the fluorescence intensity of non-transduced CTL (‘GFP^+++^’)(Figs. 1B, C, S1A, B). The percentage of CTL transduced to express the MEL5 TCR that were positive for staining with an HLA-A*0201/MART-1 ELA tetramer was only slightly higher (Fig. S1A, B).

Nevertheless, GFP^+++^ CTL could kill effectively (Fig. 1D). Cytolysis was determined as a reduction in the red area of peptide-loaded mCherry-transfected target cells adhering to the bottom of a 384-well imaging plate. A rate of 10% red area loss per hour constitutes efficient killing (Ambler et al. 2020). The cytolytic ability of GFP^+++^ CTL was rapidly declining over time of tissue culture (Fig. 1D). This was likely caused by decreasing expression of the transgenic TCR, indicated by reductions in the percentage of Vβ13.1-positive cells in the GFP^+++^ sort gate, decreased mean fluorescent intensity (MFI) of the Vβ13.1 staining, by reductions in the GFP MFI in the GFP^+++^ sort gate and a loss of blasting cell size (Fig. 1C, E-G, S1C, D). Therefore, going forward we have sorted CTL at day 7 of culture and used them the same day.

Next, we sorted CTL on day 7 of cell culture for different levels of GFP expression (Fig. 2A). Only CTL with GFP expression at least 10-fold above the fluorescence intensity of non-transduced CTL, GFP^+++^, yielded substantial cytolysis (Fig. 2B). Therefore, going forward we have used GFP^+++^ CTL. Next, we varied effector to target cell ratios in the killing assay from 1:1 to 4:1. Substantial killing occurred only at an effector to target cell ratio of 4:1 (Fig. 2C), as used going forward. In comparison, killing of neoantigen-expressing murine renal carcinoma cells by murine TCR transgenic T cells in the same imaging-based killing assay at the same high concentration of agonist peptide was already effective at an effector to target cell ratio of 1:1 (Ambler et al. 2020). Less efficient killing of the human CTL is consistent with more limited expression of the transgenic TCR in the human CTL on the background on endogenous TCR expression and/or with more limited affinity of the 1G4 and MEL5 TCRs for a tumor-associated antigen rather than a neo-antigen in the murine system as further investigated below. Human CTL were derived from blood donors such that MHC restriction of the T cells of the donor was not necessarily matched to MHC alleles expressed on the target cell lines. Moreover, the human CTL retained an endogenous TCR repertoire. Therefore, donor-to-donor variability could be expected. However, such variability was limited. Across eight donors, CTL viability after 7d of culture was above an average of 60% for each donor but one, donor 17 (Fig. 2D). Transduction efficiency for the highest GFP expression level was consistently around 20% except for donor 17 where lower viability correlated with lower transduction efficiency (Fig. 2E). Across 15 donors, there was substantial variability in killing efficiency between individual CTL cultures but not across donors (Fig. 2F). While donor-to-donor variability was moderate, we nevertheless have consistently executed experiments using cells from multiple donors because of the rare occurrence of a donor yielding CTL with substantially altered properties. Given the variability between individual CTL cultures, multiple experimental repeats were required.

**Figure 2.**
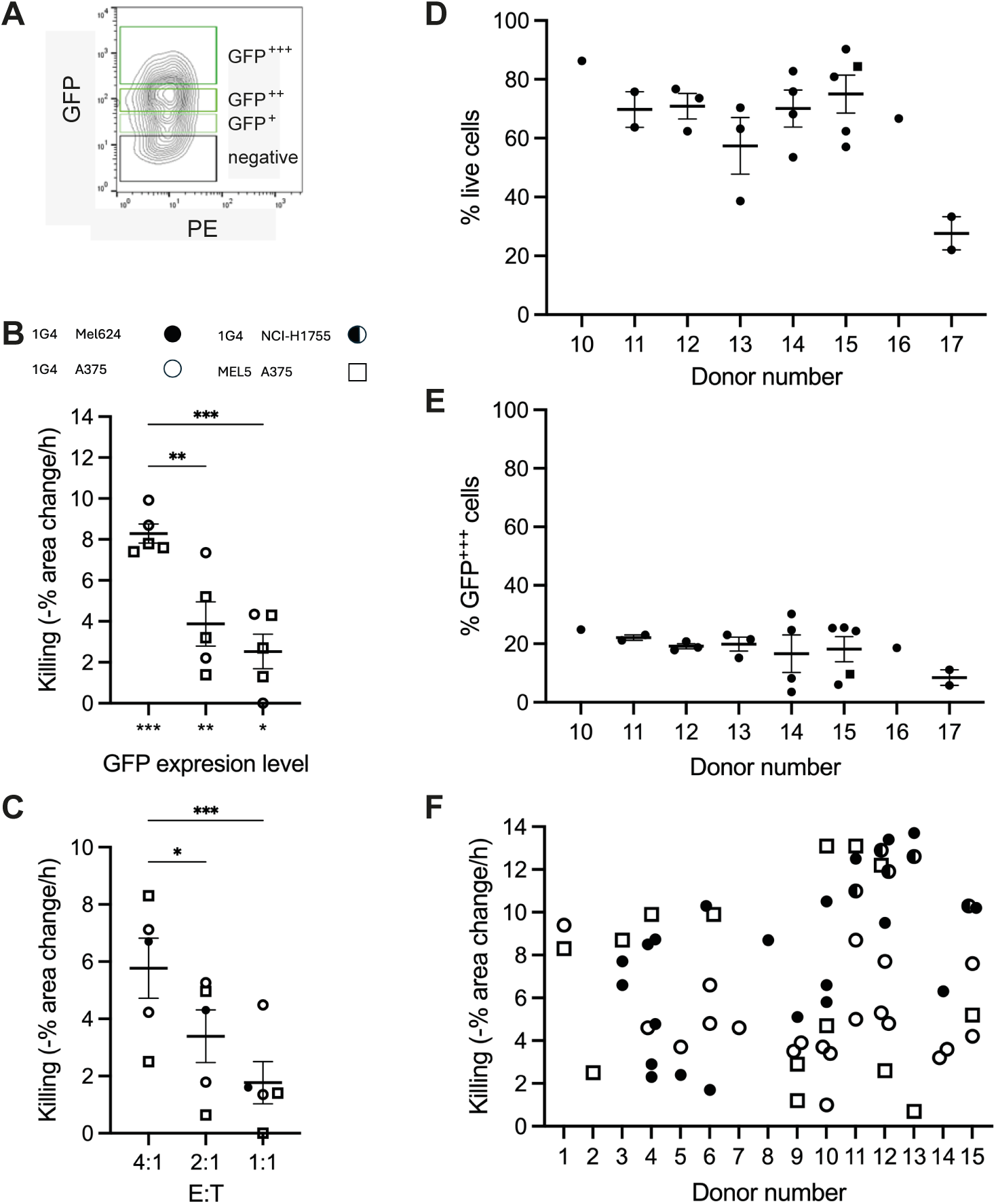
Technical parameters determining the killing of tumor target cells by primary human CTL expressing a transgenic TCR **A** Representative GFP sort windows of live CTL transduced to express the 1G4 TCR with GFP as the transduction marker. 1 of 5 independent experiments. **B** Killing of A375 tumor target cells by CTL transduced to express the 1G4 or MEL5 TCR and sorted for different levels of GFP expression as defined in A as mean ± SEM. 5 independent experiments. Statistical significance determined by paired One-way ANOVA. **C** Killing of A375 or Mel624 tumor target cells by CTL transduced to express the 1G4 or MEL5 TCR and sorted for GFP^+++^ expression at the indicated effector to target cell (E:T) ratios as mean ± SEM. 5 independent experiments. Statistical significance determined by paired One-way ANOVA. **D, E** Percentage of live cells (D) and of cells with GFP^+++^ expression (E) of total events for CTL transduced to express the 1G4 (circle) or MEL5 (square) TCR as mean ± SEM. 21 independent experiments. **F** Killing of A375, Mel624 or NCI-H1755 tumor target cells by CTL transduced to express the 1G4 or MEL5 TCR and sorted for GFP^+++^ expression by blood donor. 60 independent experiments. * p<0.05, ** p<0.01, *** p<0.001.

### Primary human CTL expressing a transgenic TCR can respond to endogenous antigen only when the antigen is highly expressed

To determine whether killing of tumor target cells by human primary CTL expressing a transgenic TCR is dependent on the recognition of antigen on the surface of the target cells by the transgenic TCR, we varied the amount of antigenic peptide on the tumor target cells. As secretion of IFNγ is a key second effector function of CTL that can be regulated differently than target cell killing, we also measured the amount of IFNγ at the end of the imaging-based cytolysis assays in the assay supernatant. First, we determined the expression of the tumor-associated antigen NY-ESO-1 in our target cell lines. Based on flow cytometry after intracellular antigen staining, all three target cell lines expressed endogenous NY-ESO-1. They did in the order of NCI-H1755 > A372 > Mel624 with 2-4-fold differences in expression between the cell lines (Fig. 3A, S3A). To enable investigations in the presence of even lower amounts of agonist peptide/MHC using 1G4 TCR-expressing CTL (1G4 CTL), we knocked down NY-ESO-1 in Mel624 cells (Fig. 3A, S3A, B). A375 cells have previously been shown to lack MART-1 (Cormier et al. 1999, Carrabba et al. 2003), Mel624 cells to express it, as confirmed here (Fig. S3C) (Cormier et al. 1999). We noted rapid recovery of NY-ESO-1 expression in the knock down cells, possibly because of the location of the NY-ESO-1 gene on a duplicated region of the x chromosome. Therefore, we used NY-ESO-1 knock down cells only for brief periods after thawing fresh aliquots.

**Figure 3.**
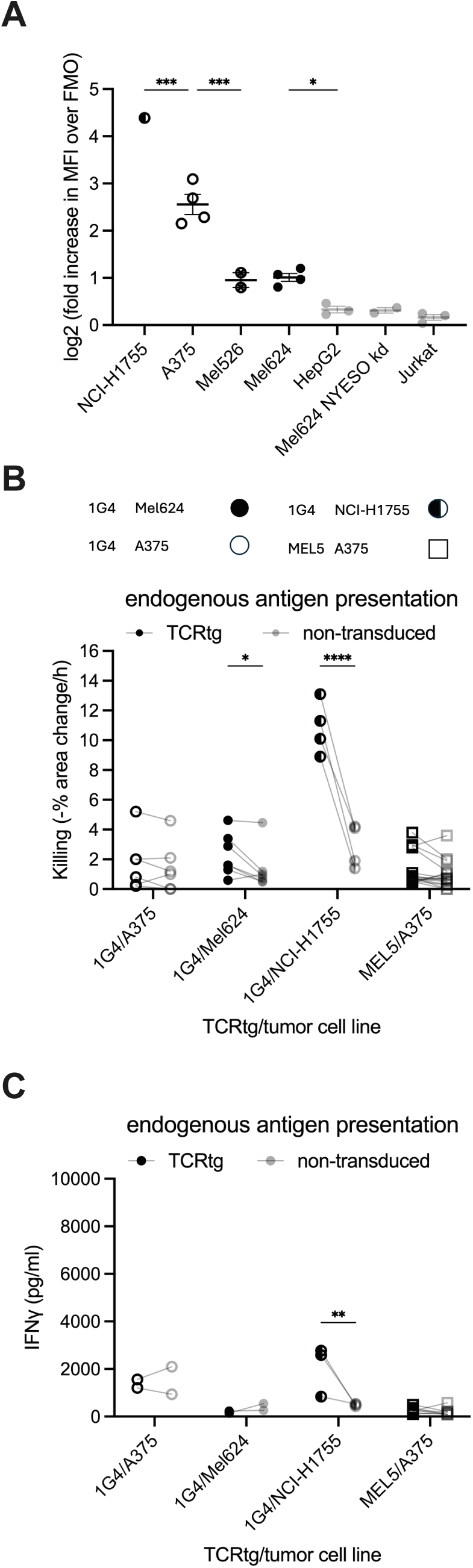
Primary human CTL expressing a transgenic TCR respond to highly expressed endogenous antigen **A** Quantification of flow cytometric NY-ESO-1 staining of indicated cells as log2 increase in MFI over fluorescence minus one (FMO) as mean ± SEM. Representative flow cytometry data in Fig. S3A. 1 to 4 independent experiments. Statistical significance determined by One-way ANOVA. **B** Killing of A375, Mel624 or NCI-H1755 tumor target cells by 1G4 or MEL5 CTL in comparison to CTL with an endogenous TCR repertoire (‘non-transduced’). 33 independent experiments. Statistical significance determined by paired Two-way ANOVA. **C** IFNγ amounts in supernatants of A375, Mel624 or NCI-H1755 tumor target cells after 16h of interaction with 1G4 or MEL5 CTL in comparison to CTL with an endogenous TCR repertoire (‘non-transduced’). 16 independent experiments. Statistical significance determined by paired Two-way ANOVA. * p<0.05, ** p<0.01, *** p<0.001, **** p<0.0001.

To determine whether expression of a transgenic TCR in human primary CTL could confer antigen specificity in response to endogenous antigen presentation, we compared the ability of CTL transduced to express a transgenic TCR to kill tumor target cells in the absence of exogenous agonist peptide to that of primary human CTL expressing only an endogenous TCR repertoire (Fig. 3B). 1G4 CTL effectively killed NCI-H1755 target cells and moderately but significantly (p<0.05) Mel624 cells. Recognition of antigen expressed by NCI-H1755 was also sufficient to trigger some IFNγ secretion (Fig. 3C). While CTL expressing the 1G4 or MEL5 TCR could kill A375 cells to a moderate extent in some experimental repeats, such killing was indistinguishable from that by CTL not expressing a transgenic TCR and not associated with antigen-specific IFNγ secretion (Fig. 3B, C). Endogenous antigen thus could only trigger an antigen-specific functional response of the TCR expressing CTL when it was highly expressed.

### Agonist peptide amount and affinity and the TCR expression level control the effector function of human CTL expressing a transgenic TCR

To determine whether higher concentrations of agonist peptide can trigger TCR-mediated activation of the human CTL, we incubated the target cell lines with a high concentration, 2µg/ml, of agonist peptide. The additional agonist peptide triggered significant (p<0.001) increases in target cell killing in all TCR transgene/target cell combinations except for the combination of human 1G4 CTL interacting with NCI-H1755 cells where killing was already efficient in the absence of exogenous peptide (Fig. 4A). Highly efficient killing was seen in at least a few experimental repeats in all TCR transgene/target cell combinations. In addition, substantial increases in IFNγ secretion were observed (Fig. 4B). The interaction of CTL transduced to express the MEL TCR (MEL5 CTL) with target cells lacking MART-1, A375, as corroborated in the interaction of 1G4 CTL with target cells with a very low expression of NY-ESO-1, Mel624 NY-ESO-1 kd, established a low cytolysis background, around less than 2% red area change per hour, and a low background of IFNγ secretion, around less than 500pg/ml (Fig. 4A, B). Interaction of MEL5 CTL with the MART-1-negative A375 cells occasionally showed higher killing, up to 4% red area change per hour, but consistently low IFNγ secretion, less than 500pg/ml (Fig. 4A, B). Thus, a background of CTL activation not driven by a transgenic TCR recognizing its cognate peptide/MHC complex is low but varies between TCRs and/or cell lines, as further characterized below. Using a 10-fold peptide dose titration for a subset of the 1G4 CTL data, an agonist peptide concentration of 2ng/ml was saturating for cytolysis and IFNγ secretion (Fig. 4C, D). We next verified that killing and IFNγ secretion in the presence of 2µg/ml agonist peptide were TCR-dependent through comparison with non-transduced CTL. Killing and IFNγ secretion were consistently enhanced upon expression of a transgenic TCR (Fig. 4E, F), supporting antigen-specificity of CTL effector function.

**Figure 4.**
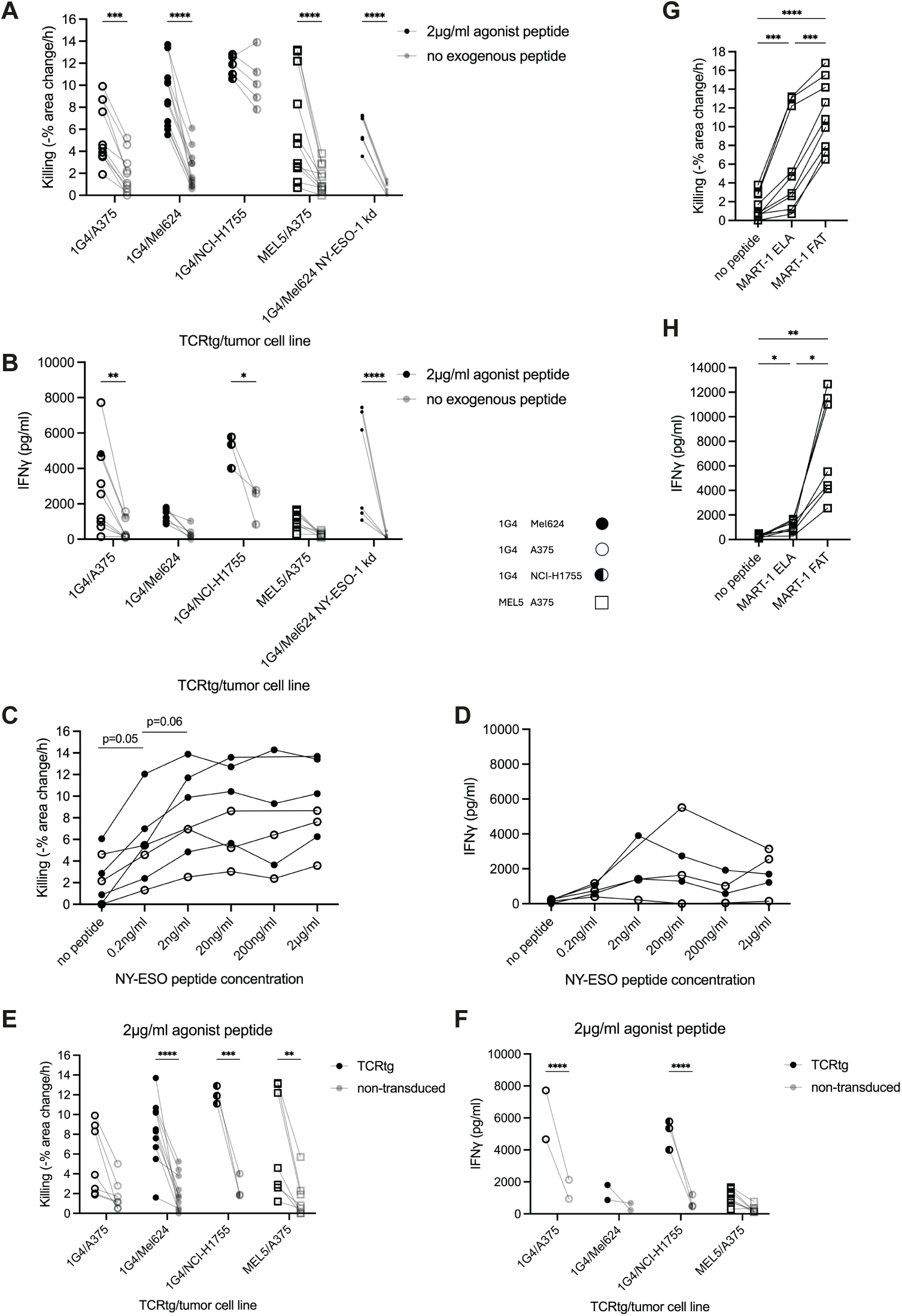
Agonist peptide amount and affinity determine the function of primary human CTL expressing a transgenic TCR **A** Killing of A375, Mel624, Mel624 NY-ESO-1 kd or NCI-H1755 tumor target cells incubated with or without 2µg/ml NY-ESO-1 or ELA MART-1 agonist peptide by 1G4 or MEL5 CTL. 42 independent experiments. Statistical significance determined by paired Two-way ANOVA. **B** IFNγ amounts in supernatants of A375, Mel624, Mel624 NY-ESO-1 ko or NCI-H1755 tumor target cells incubated with or without 2µg/ml NY-ESO-1 or ELA MART-1 agonist peptide after 16h interaction with 1G4 or MEL5 CTL. 31 independent experiments. Statistical significance determined by paired Two-way ANOVA. **C** Killing of A375 or Mel624 tumor target cells incubated with the indicated concentration of NY-ESO-1 agonist peptide by 1G4 CTL. 7 independent experiments. Statistical significance determined by paired Two-way ANOVA. **D** IFNγ amounts in supernatants of A375 or Mel624 tumor target cells incubated with the indicated concentration of NY-ESO-1 agonist peptide after 16h interaction with 1G4 CTL. 5 independent experiments. **E** Killing of A375, Mel624 or NCI-H1755 tumor target cells incubated with 2µg/ml NY-ESO-1 or ELA MART-1 agonist peptide by 1G4 or MEL5 CTL in comparison to CTL with an endogenous TCR repertoire (‘non-transduced’). 27 independent experiments. Statistical significance determined by paired Two-way ANOVA. **F** IFNγ amounts in supernatants of A375, Mel624 or NCI-H1755 tumor target cells incubated with 2µg/ml NY-ESO-1 or ELA MART-1 agonist peptide after 16h interaction with 1G4 or MEL5 CTL in comparison to CTL with an endogenous TCR repertoire (‘non-transduced’). 14 independent experiments. Statistical significance determined by paired Two-way ANOVA. **G** Killing of A375 tumor target cells incubated with or without 2µg/ml ELA or FAT MART-1 agonist peptide by MEL5 CTL. 9 independent experiments. Statistical significance determined by paired One-way ANOVA. **H** IFNγ amounts in supernatants of A375 tumor target cells incubated with or without 2µg/ml ELA or FAT MART-1 agonist peptide after 16h interaction with MEL5 CTL. 7 independent experiments. Statistical significance determined by paired One-way ANOVA. * p<0.05, ** p<0.01, *** p<0.001, **** p<0.0001.

We noted that IFNγ secretion in MEL5 CTL was only moderately triggered by A375 cells incubated with the high concentration of exogenous antigenic peptide (Fig. 4F). To determine whether such moderate T cell activation could be a consequence of a limited affinity of the TCR for the tumor-associated antigen-derived peptide/MHC complex, we used the FAT variant of the MART-126-35 peptide that yields a higher affinity of the MEL5 TCR for the peptide/MHC complex with a Kd of 3µM as compared to 17µM for the ELA variant (Ekeruche-Makinde et al. 2012).

Incubating A375 cells with 2µg/ml of the FAT peptide trigger a 1.8-fold increase in cytolysis and a 7.1-fold increase in IFNγ secretion in comparison to the ELA variant of the MART-1 peptide (Fig. 4G, H). As only moderate TCR affinities for cognate peptide/MHC complexes are common in anti-tumor immunity, we kept mostly using them here.

Transgenic TCRs expressed in parallel with endogenous ones can be stabilized using an additional disulfide bridge or by exchanging the human constant domains with the murine ones (Cohen et al. 2006, Cohen et al. 2007). We used ELA and FAT peptides and A375 cells to compare CTL function upon MEL5 TCR stabilization through a disulfide bridge versus the use of murine constant domains. Primary human CTL were lentivirally transduced to express the MEL5 TCR variants (Fig. S4A) and sorted for equal expression of GFP as the transduction marker.

Killing of A375 target cells was indistinguishable in the absence of agonist peptide and in the presence of 2µg/ml of the high affinity FAT peptide. Killing in response to 2µg/ml of the moderate affinity ELA peptide was moderately more effective in CTL expressing the MEL5 TCR with the murine constant domains (Fig. S4B). In contrast to the minor effects on killing, IFNγ secretion was 39-fold and 4-fold enhanced upon MEL5 stabilization with the murine constant domains in response to 2µg/ml of the ELA and FAT peptides, respectively (Fig. S4C). A likely explanation for the enhanced function of CTL expressing the murine constant domain-stabilized MEL5 TCR was substantially increased MEL5 TCR expression as determined with tetramer staining (Fig. S4D). As TCR stabilization using only an additional disulfide bridge is common in therapeutic applications, we kept using it here.

In summary, activation of primary human CTL expressing a recombinant TCR by interaction with tumor target cells could be effective yet was more often limited by low expression of tumor-associated antigens in the target cells, the only moderate affinity of the TCR for the peptide/MHC complexes and limited TCR expression when using an additional disulfide bridge to stabilize the recombinant TCR. These limitations were more severe in cytokine secretion than target cell killing and could be overcome by incubation of tumor cells with exogenous agonist peptide, by using higher affinity peptide variants and by stabilizing the TCR with murine constant domains. Nevertheless, one of four combinations of human CTL expressing a disulfide bridge- stabilized recombinant TCR and tumor target cells tested, the interaction of 1G4 TCR-expressing CTL with NCI-H1755 target cells, allowed for the investigation of CTL activation even in response to endogenous antigen presentation.

### Primary human CTL display graded subcellular organization in response to stimuli ranging from no antigen to high concentrations of high affinity antigen

Even in the absence of antigen or the presence of very low amounts thereof, a low background of CTL function could be observed depending on the TCR tumor cell combination (Fig. 4A, B). These data suggest that CTL respond across a wide range of stimuli strengths. The ability of CTL to form tight cell couples and maintain them in an F-actin-dependent fashion is a key cellular mechanism underpinning target cell killing and IFNγ secretion (Ritter et al. 2015, Ambler et al. 2020). To further characterize CTL activation across stimulus strengths, we investigated two CTL tumor target cell combinations that together cover the entire stimulus range established in the functional assays, the interaction of murine constant domain-stabilized MEL5 TCR-expressing CTL (MEL5mC CTL) with A375 cells in response to a high concentration of the high affinity FAT peptide or no antigen (Fig. S3C, 4A) and the interaction of 1G4 CTL with NY-ESO-1 wild type, plus/minus additional agonist peptide, or NY-ESO-1 knock down Mel624 cells (Figs. 3A, 4A), as described sequentially here.

The conversion of an initial contact of a CTL with a target cell into a tight cell couple characterized by an interface as wide as the CTL is an active, F-actin-dependent process and thus indicative of the strength of the signal perceived by the CTL during the initial contact. As T cells can only form tight cell couples upon target cell contact with the leading edge (Negulescu et al. 1996), cell coupling frequencies of 50% and above are indicative of effective initial target cell recognition. MEL5mC CTL formed tight cell couples in 32±5% of the initial contacts with A375 target cells in the absence of antigen, less than upon addition of 2µg/ml of the FAT agonist peptide but nevertheless indicative of effective target cell recognition in the absence of antigen (Fig. 5A, B). Effective killing is associated with the maintenance of a polarized CTL target cell couple (Fig. 5C-G). Two measurements of cell couple stability are the delayed formation of lamellae pointing away from the interface and lack of translocation of the CTL over the target cell surface (Ambler et al. 2020). In the absence of agonist peptide, cell couples of MEL5mC

**Figure 5.**
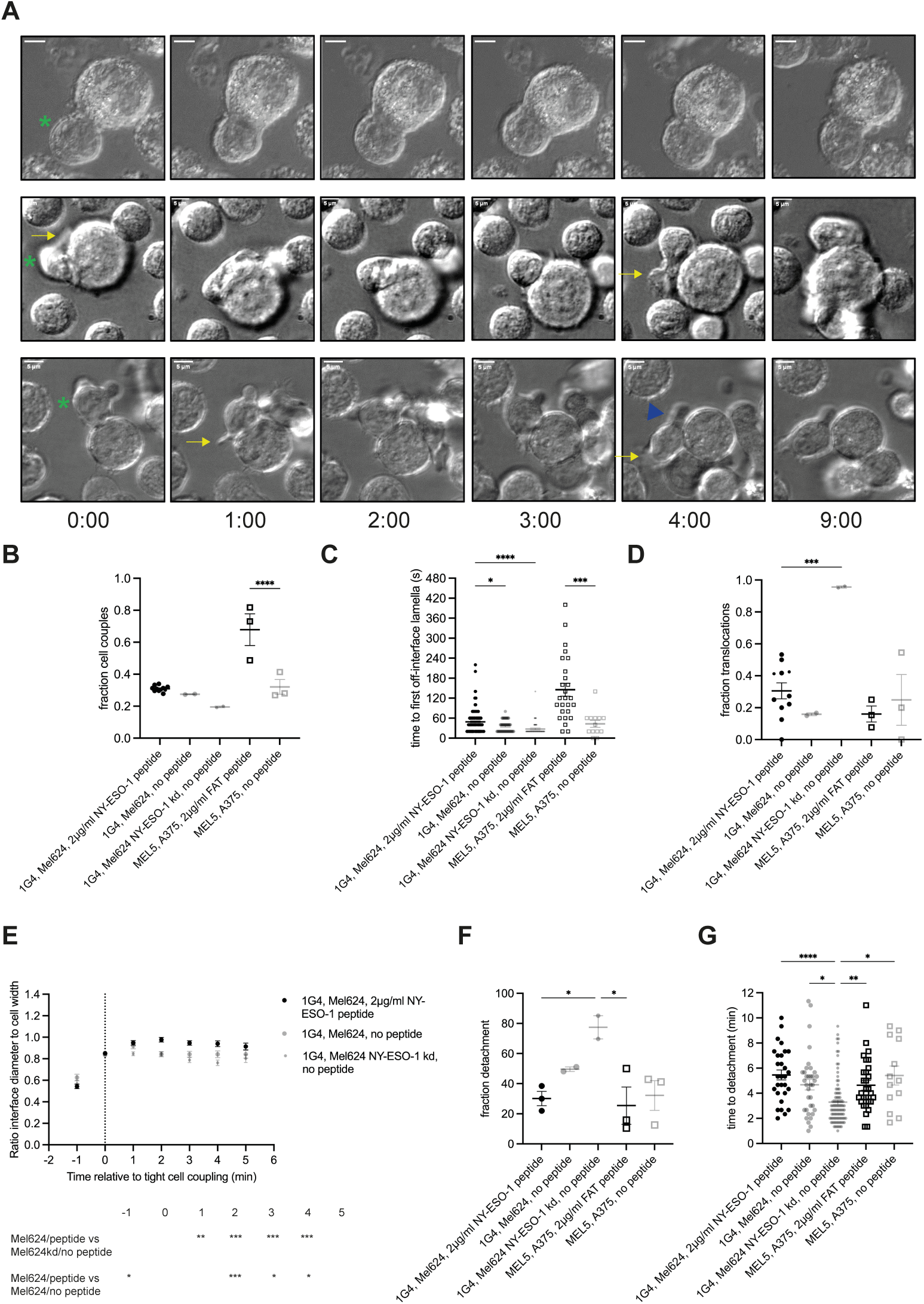
CTL morphology is regulated by stimulus strength in a graded fashion **A** Representative bright field imaging data of the interaction between 1G4 CTL and wild type (top row) and NY-ESO-1 knock down (middle and bottom rows) Mel624 cells in the absence of exogenous agonist peptide. Time relative to tight cell couple formation in minutes is given below the panels. The CTL is denoted with a green asterisk. The top row shows a stable cell couple, the middle row translocation and the bottom row detachment. Off-interface lamellae are indicated with a yellow arrow, residual attachment through the uropod with a blue triangle. 1 representative experiment of 2. Scale bar = 5µm. **B-G** Characterization of cell morphology in the interaction of 1G4 or MEL5 CTL with Mel624, wild type (large symbol) or NY-ESO-1 knock down (small symbol), or A375 cells, respectively, in the presence of the given amount of the indicated agonist peptide as mean ± SEM. The MEL5 CTL used in these imaging experiments also express a chimeric costimulatory receptor in the absence of any ligand. B Fraction of CTL converting a target cell contact into a tight cell couple. Each symbol is an imaging run. Small symbols in the MEL624 + peptide condition indicate a subset of the 1G4 CTL data matching the Mel624 and Mel624 NY-ESO-1 kd conditions. C Time from tight cell couple formation to first off- interface lamella. Each symbol is a cell couple. D Fraction of CTL with a translocation. Small symbols in the MEL624 + peptide condition indicate a subset of the 1G4 CTL data matching the Mel624 and Mel624 NY-ESO-1 kd conditions. E Interface diameter relative to the CTL width of 1G4 CTL. Single cell data in Fig. S5C. MEL5 CTL data in Fig. S5A, B. F Fraction of CTL with detachment. Each symbol is an imaging run. G Time from tight cell couple formation to detachment. Each symbol is a cell couple. 2-8 independent experiments. Statistical significance determined by One-way ANOVA (B, D, F), Two-way ANOVA (E) and Kruskall-Wallis test (C, G). * p<0.05, ** p<0.01, *** p<0.001, **** p<0.0001.

CTL with A375 target cells formed off-interface lamellae more rapidly and showed a higher frequency of translocations (Fig. 5C, D), indicative of inefficient cell couple maintenance. Consistent with less efficient formation of cell couples in the absence of agonist peptide, interface diameters were smaller upon initial cell coupling (Fig. S5A, B). A complete loss of polarization towards the interface, ‘detachment’, was defined as consistent leading edge lamellae at the opposite side of the CTL target cell interface (Fig. 5A). Breaking of the CTL target cell membrane contact occurred substantially later, if it could be observed on the time scale of the imaging experiments at all. Such detachment occurred in a minority of cell couples with no difference in frequency and time to detachment in the presence versus absence of 2µg/ml FAT agonist peptide (Fig. 5F, G). Cell couple formation is associated with overall F-actin accumulation as imaged using F-tractin-GFP (Yi et al. 2012) at the cellular interface. The rapid clearing of F-actin at the interface center (Fig. 6A) is required for effective progression of the cell couple to cytolysis (Ritter et al. 2015, Ambler et al. 2020). Central F-actin clearance was absent in MEL5mC CTL in the absence of antigen (Fig. 6B, S6A), while overall interface F-actin accumulation was not impaired (Fig. 6C, S6B). Together these data suggest that MEL5mC CTL perceive a substantial stimulus when contacting A375 target cells in the absence of antigen, triggering effective cell coupling and F-actin recruitment to the cellular interface. However, the stimulus was too weak to allow effective cell couple maintenance in preventing immediate off-interface lamellae and translocations, full generation of a wide interface and central F-actin clearance, consistent with target cell killing at or just above background (Fig. 4A) and the lack of IFNγ secretion (Fig. 4B).

**Figure 6.**
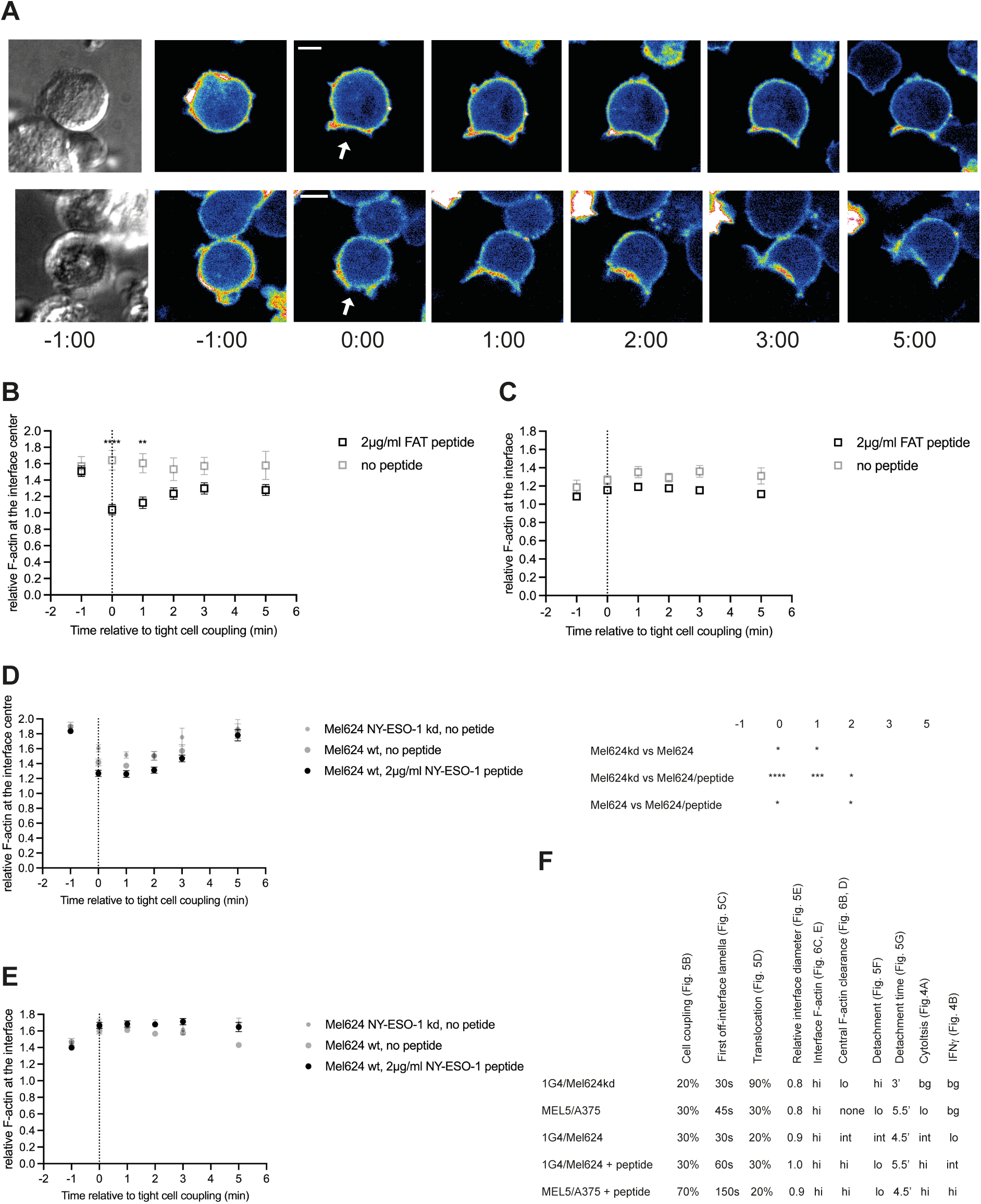
Only effective cytolysis is associated with F-actin clearance at the center of the CTL target cell interface **A** Representative spinning disk confocal imaging data of 1G4 TCR and F-tractin-GFP expressing CTL with Mel624 cells, incubated with 2 µg/ml NY-ESO-1 peptide (top) or not (bottom). At the left a differential interference contrast image is shown at the first time point. It is followed by midplane images of F-tractin-GFP distributions at the indicated time points relative to tight cell coupling. The cellular interface is indicated with a white arrow at the time of tight cell coupling. The top row shows effective F-actin clearance at the interface center with noticeable F-actin accumulation at the interface edge, the bottom row lack thereof. Scale bar = 5µm **B, C** F-actin accumulation at the interface between CTL expressing the MEL5 TCR with F-tractin- GFP and A375 cells in the presence of the indicated amounts of agonist peptide relative to F- actin in the entire cell and to the time of tight cell coupling as mean ± SEM. Pooled data from 3 independent experiments. B F-actin accumulation at the central third of the interface. Single cell data in Fig. S6A. Statistical significance determined by Two-way ANOVA. **C** F-actin accumulation at the entire interface. Single cell data in Fig. S6B. **D, E** F-actin accumulation at the interface between CTL expressing the 1G4 TCR with F-tractin-GFP and Mel624 cells, wild type or NY-ESO-1 kd, in the presence of the indicated amounts of agonist peptide relative to F- actin in the entire cell and to the time of tight cell coupling as mean ± SEM. Pooled data from 2- 3 independent experiments. D F-actin accumulation at the central third of the interface. Single cell data in Fig. S6C. Statistical significance determined by Two-way ANOVA and displayed in the table to the right. E F-actin accumulation at the entire interface. Single cell data in Fig. S6D. **F** Comparison of CTL morphological features with effector function across an increasing range of CTL stimulus strength with figures containing raw data indicated. ‘bg’ is background. * p<0.05, ** p<0.01, *** p<0.001, **** p<0.0001.

In the interaction of 1G4 CTL with Mel624 target cells stimulus strength was varied from a high concentration of exogenous NY-ESO-1 agonist peptide to endogenous antigen expression to knock down of NY-ESO-1 expression. Cell couple formation and maintenance were highly inefficient in the interaction of 1G4 CTL with NY-ESO-1 knock down Mel624 target cells. The cell coupling frequency was low at 19%, off-interface lamellae were instantaneous, translocation occurred in virtually all cell couples and the CTL target cell interface was narrower (Fig. 5B-E, S5C). 76±9% of 1G4 CTL cell couples with NY-ESO-1 knock down Mel624 target cells detached with a significantly (p<0.05) shortened time to detachment of 3.3±0.2min (Fig. 5F, G). While F-actin was effectively recruited to the cellular interface upon cell coupling, central F-actin clearance was inefficient (Fig. 6D, E, S6C, D). Cell couples between 1G4 CTL and NY-ESO-1 knock down Mel624 target cells, while they still formed, thus were highly unstable. Whether the marginal residual cell coupling was still dependent on 1G4 TCR recognition of the NY-ESO-1 peptide/MHC complex was not resolved. Cell coupling of 1G4 CTL to wild type Mel624 cells upon endogenous antigen expression was more effective than that triggered by NY-ESO-1 knock down Mel624 cells. However, it still led to less stable cell couples than seen in the presence of a high concentration of agonist peptide, consistent with limited target cell killing and lack of IFNγ secretion (Fig. 4A, B). While cell coupling was moderately efficient, translocations were almost completely prevented and interfaces were wider (Fig. 5B, D, E), first off-interface lamellae occurred still significantly (p<0.05) faster than in the presence of a high concentration of agonist peptide (Fig. 5C). Central F-actin clearance showed an intermediate phenotype (Fig. 6D, S6C). In combination with the MEL5mC CTL data the 1G4 CTL data establish that the subcellular organization of CTL in target cell couples gradually change from extremely unstable to fully stabilized across a wide range of stimuli strengths. Such subcellular CTL organization, more than cell couple formation per se, was related to CTL effector function (Fig. 6F).

Together the 1G4 CTL and MEL5 CTL data also establish that what constitutes a limiting stimulus varies with the TCR target cell combination. The level of cell coupling and maintenance in one CTL target cell combination in the absence of antigen, MEL5mC CTL and A375 cells here, became as high as the level of cell coupling and maintenance in another CTL target cell combination in the presence of endogenous antigen presentation, 1G4 CTL and Mel624 cells here.

### Interaction of primary human CTL expressing a transgenic TCR with tumor cell spheroids induces CTL suppression

In previous work, we have established that the interaction of in vitro primed murine TCR transgenic CTL with tumor cell spheroids in the presence of the cognate antigen triggers a phenotype in the spheroid-infiltrating lymphocytes (‘SIL’) that closely resembled that of suppressed tumor-infiltrating lymphocytes (Ambler et al. 2020). Key features of this phenotype are impaired effector function, upregulation of inhibitory receptor expression, a severe calcium signaling defect and a reduced ability to maintain a fully polarized cell couple with target cells (Ambler et al. 2020). Based on this work, we determined whether incubation of human primary CTL expressing a transgenic TCR with tumor cell spheroids could trigger similar suppression.

To allow effective suppression of CTL, tumor cell lines must be able to form spheroids and such spheroids must be sufficiently stable to allow the isolation of spheroid-interacting CTL for subsequent analysis. Spheroid generation needed to be experimentally optimized for each tumor cell line. A375 and Mel624 spheroids were grown from single cell suspensions in Matrigel for 12 days to an average size of around 200-500µm (Fig. S7A, B) forming spherical structures (Fig. S7C). Matrigel was dissolved and spheroids were washed, a procedure required in the isolation of spheroid-infiltrating CTL as described in the next paragraph. While Mel624 spheroids consistently remained intact throughout this washing procedure, A375 spheroids were falling apart in a subset of experiments. The impaired ability of A375 spheroids to form a smooth surface (Fig. S7D) may be indicative of weaker cell-to-cell adhesion, as consistent with reduced spheroid stability. NCI-H1755 cells did not form spheroids well in Matrigel. Instead, NCI-H1755 spheroids needed to be grown in agarose-coated round bottom 96 well plates in the presence of a small concentration of Matrigel, 2.5%, as to be characterized in detail elsewhere.

To induce CTL suppression, GFP^+++^ 1G4 CTL were mixed with spheroids that had been incubated with 2µg/ml NY-ESO-1 agonist peptide for 1h and both cell types were embedded together in Matrigel and cultured for 16h in the absence of IL-2. Live cell imaging of the interaction of 1G4 CTL with spheroids confirmed rapid and deep infiltration of CTL into the spheroids (Fig. S7E-G). Matrigel was dissolved, spheroids were washed to remove unbound CTL and then mechanically disaggregated. Spheroid-infiltrating CTL (spheroid-infiltrating lymphocytes, ‘SIL’) were purified by fluorescence-activated cell sorting for GFP^+++^ expression. Comparing 1G4 SIL to CTL that were kept in tissue culture in the presence of IL-2, killing was significantly (p<0.05) diminished by on average 40%, IFNγ secretion by 79%, being undetectable in most experimental repeats (Fig. 7A, B). Overnight incubation of human primary CTL expressing a transgenic TCR with tumor cell spheroids presenting the cognate agonist peptide thus induced functional suppression in the CTL. Next, we investigated whether spheroids must be incubated with a high concentration of exogenous agonist peptide to induce CTL suppression or whether endogenous antigen presentation suffices. Using the interaction of 1G4 CTL with Mel624 spheroids, the ability of SIL to kill tumor target cells and secrete IFNγ was equally suppressed whether 2µg/ml exogenous NY-ESO-1 peptide was present in the spheroids or not (Fig. 7C, D). As most data have been acquired using SIL induction in the presence of exogenous peptide, we continue doing so here.

**Figure 7.**
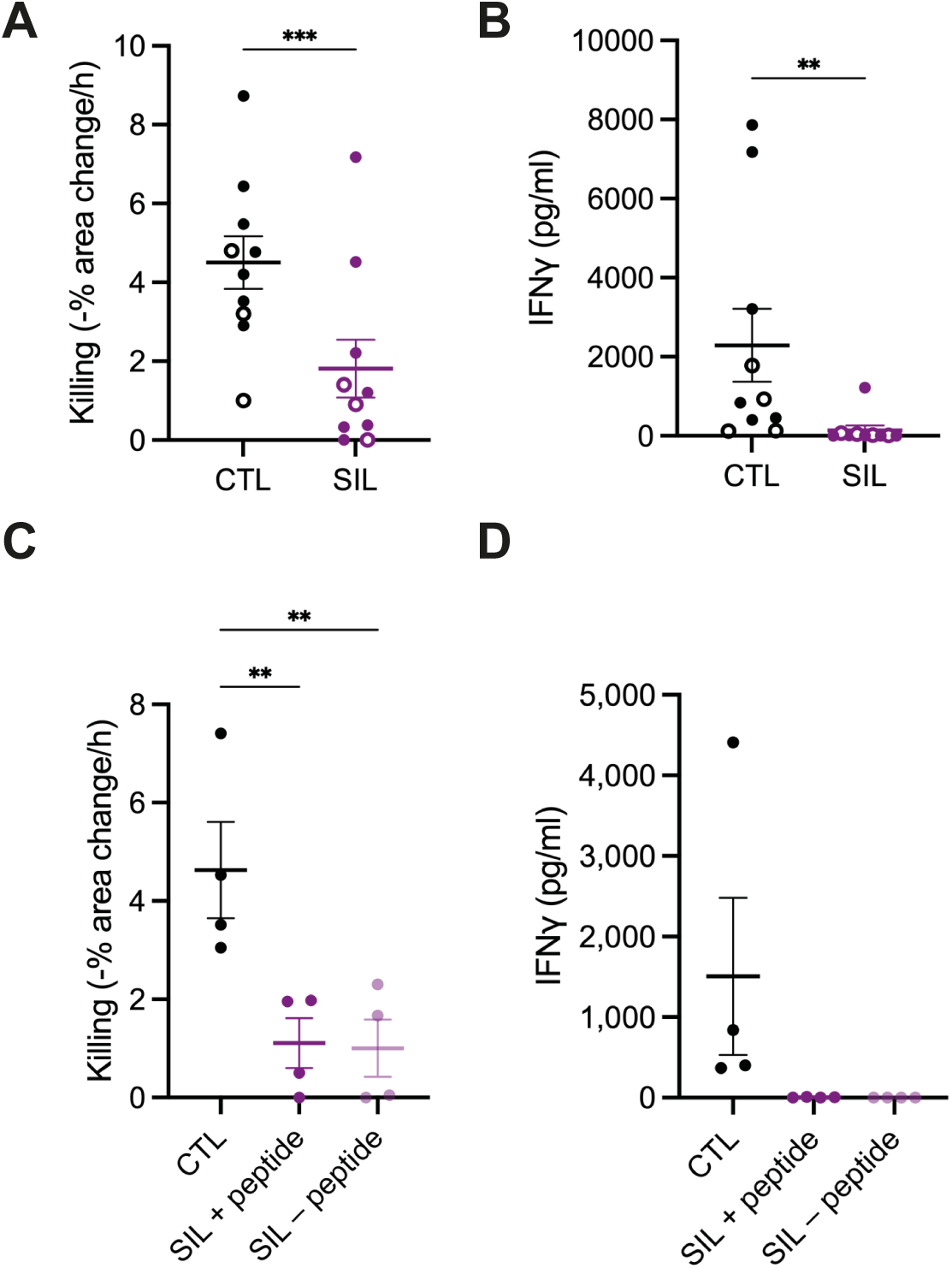
CTL interaction with spheroids induces a CTL suppression **A, B** Killing of A375 (open symbols) or Mel624 (closed symbols) tumor target cells incubated with 2µg/ml NY-ESO-1 agonist peptide by 1G4 CTL or SIL and IFNγ amounts in supernatants after 16h interaction as mean ± SEM. 10 independent experiments. Statistical significance determined by paired Student’s t-test. A Killing, B IFNγ amounts **C, D** Killing of Mel624 tumor target cells incubated with 2µg/ml NY-ESO-1 agonist peptide by 1G4 CTL or SIL with Mel624 spheroids used in the SIL generation incubated with 2µg/ml of exogenous NY-ESO-1 agonist peptide (‘SIL + peptide’) or not (‘SIL – peptide’) and IFNγ amounts in supernatants after 16h interaction as mean ± SEM. 4 independent experiments. Statistical significance determined by paired One-way ANOVA. C Killing, D IFNγ amounts. ** p<0.01, *** p<0.001.

### Suppressed CTL express high levels of exhaustion markers

To characterize the phenotype of 1G4 CTL and 1G4 SIL, we used a 21-antibody panel with parallel detection of all antibodies on a flow cytometer equipped with a spectral analyzer. For comparison, unstimulated PBMC, PBMC stimulated with anti-CD3/CD28 beads for 72h and blood and tumor-infiltrating lymphocytes from melanoma patients were also stained (Fig. 8A-D). For analysis, all populations were gated for CD45^+^CD14^−^CD19^−^CD3^+^CD8^+^ cells (Fig. S8A). 1G4 CTL were highly activated as indicated by high expression of CD25 (Fig. 8A, C) with expression of multiple inhibitory receptors, in particular LAG-3 and TIM-3, significantly (p<0.001) elevated in comparison to unstimulated PBMC (Fig. 8A, D, S8B). Such expression was also elevated in comparison to PBMC stimulated for three days, suggesting a partial induction of exhaustion in the 1G4 CTL. 1G4 CTL in comparison to stimulated PBMC displayed a significant (p<0.01) loss of CD28 expression (Fig. 8A) as associated with the induction of senescence (Huff et al. 2019), loss of responsiveness to persistent antigen (Trendel et al. 2021) and impaired self-renewal (Humblin et al. 2023). To investigate whether continuous stimulation with anti-CD3/CD28 beads over 7d in the generation of 1G4 CTL could have contributed to the partially exhausted phenotype, we set up 1G4 CTL with bead stimulation only for the first or first three days, keeping the CTL in IL-2 medium for the remainder of the culture period. Shorter stimulation led to reduced cytolytic capability (Fig. S8C), no difference in IFNγ secretion (Fig. S8D) and lower expression of PD-1 and Ki67 (Fig. S8E), consistent with a less active phenotype. Persistent 1G4 CTL stimulation thus was required for the efficient induction of cytolytic function at the expense of partial exhaustion through increased expression of inhibitory receptors.

**Figure 8.**
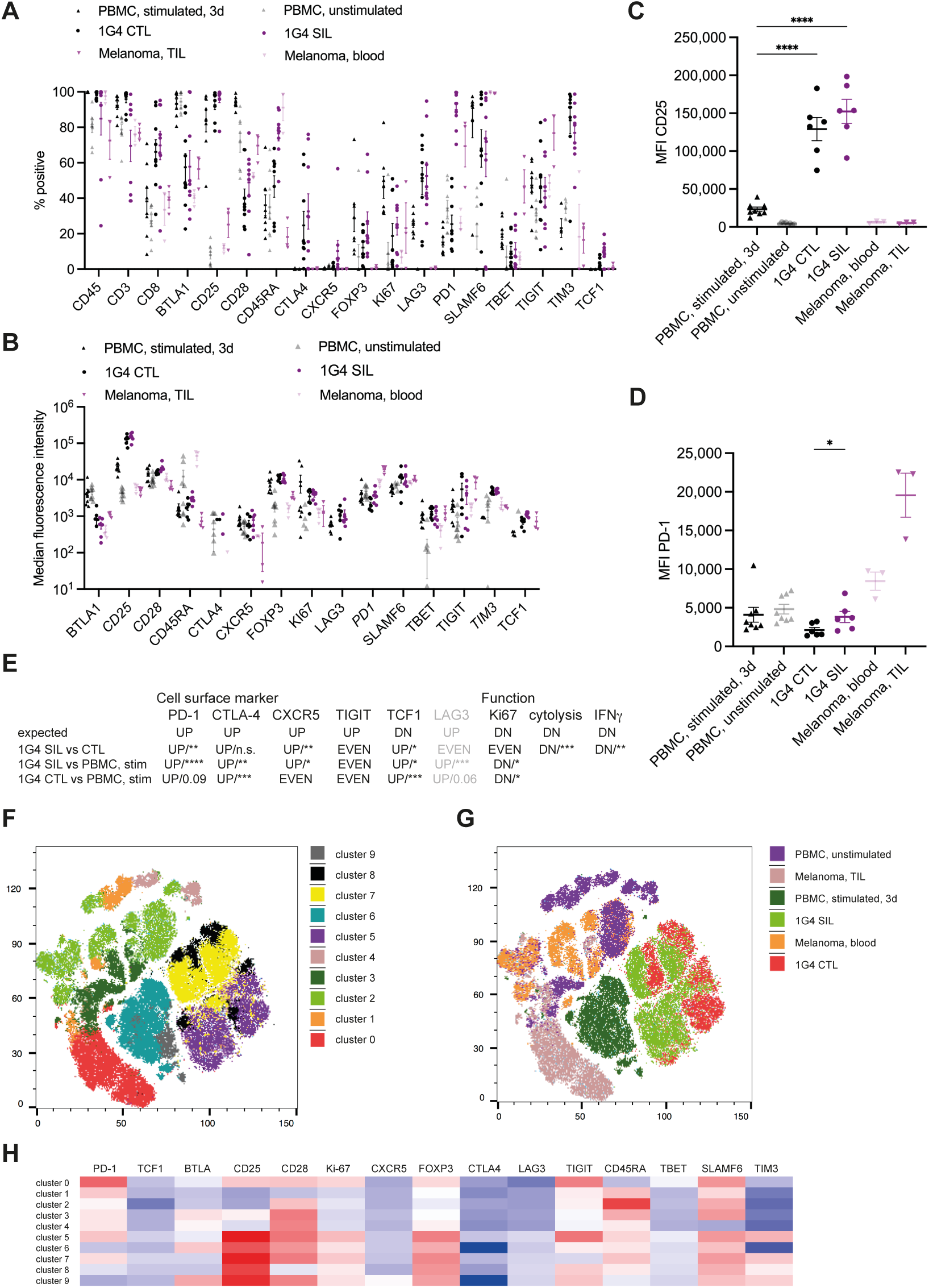
CTL interaction with spheroids induces an exhausted CTL phenotype **A-D** Percentage of CD14^−^CD19^−^CD45^+^CD3^+^CD8^+^ cells positive for the indicated markers and median fluorescence intensity as determined by flow cytometry for the indicated six experimental conditions as mean ± SEM. 3-9 independent experiments. Statistical significance determined by One-way ANOVA for individual markers. A Percentage cells positive. B Median fluorescence intensity. As a default, the MFI was determined on the entire CD8^+^ T cell population. If marker positive and negative populations could be unambiguously separated, the MFI was determined for only the positive population, as indicating by an italicized marker name. C Median fluorescence intensity of CD25 expression for the fraction of cells gated as positive from B. D Median fluorescence intensity of PD-1 expression for the fraction of cells gated as positive from B. Statistical significance of the difference between 1G4 CTL and SIL determined by paired Student’s t-test. **E** Changes in cell surface marker expression (data from A) and CTL function (data from Fig. 7A, B, 8A) as ‘expected’ for exhausted CD8^+^ T cells in comparison to naïve, effector and memory cells and changes found in the comparison of the indicated cell populations. Primary exhaustion markers in black, secondary markers in grey. Statistical significance determined by Mann-Whitney u-test in the pairwise comparisons. **F-H** Cluster analysis of the same data as in A as, F a t-SNE blot of clusters identified, G overlay of the six experimental conditions and H expression of the indicated markers. * p<0.05, **** p<0.0001.

To characterize the phenotype of the 1G4 SIL, we compared changes in cell surface marker expression and CTL function to those expected in exhausted CD8^+^ TIL. Based on an integrated analysis of scRNAseq, epigenomics and mass cytometry data (Winkler et al. 2019, O’Boyle et al. 2020) 11 primary and 16 secondary cell surface markers for the distinction of exhausted from naïve, memory and effector CD8^+^ T cells were suggested. Five of the primary and one secondary marker were part of our antibody panel (Fig. 8E). In addition, we have analyzed T cell proliferation, cytolysis and IFNγ secretion as functional markers. A comparison between PBMC stimulated for three days, 1G4 CTL and 1G4 SIL suggests a graded progression towards exhaustion. Five of seven flow cytometry-determined markers in 1G4 SIL showed an expected and significant (p<0.05) change in comparison to three-day stimulated CD8^+^ T cells, five of nine flow cytometry and functional markers in comparison to 1G4 CTL (Fig. 8E). Two of the four markers that did not change as expected, LAG-3 and Ki67, were already up/downregulated in the 1G4 CTL compared to the three-day stimulated PMBC. The percentage of CD8^+^ T cells expressing PD-1 as a prominent activation and exhaustion marker was significantly (p<0.0001) upregulated in 1G4 SIL, reaching levels comparable to melanoma patient TIL (Fig. 8A, S8B), as corroborated by an analysis of median intensity (Fig 8D, S8B). Thus, seven of the nine markers showed the expected progression to exhaustion, consistent with the suggestion that the 1G4 CTL have a partially and the 1G4 SIL an extensively exhausted phenotype that matches in vivo data well. The two exceptions were TIGIT where the fraction of cells expressing it did not change and TCF-1 that was upregulated in 1G4 CTL and SIL, albeit at a very low level, as further discussed below. To corroborate the single marker analysis, the flow cytometry data across all populations were clustered (Fig. 8F-H). Unstimulated PBMC and melanoma patient blood samples were well separated from the stimulated T cell populations (Fig. 8G). Clustering confirmed similarity between 1G4 CTL and SIL as both were distributed across clusters 5, 7, and 8 (Fig. 8F, G). Cluster 8 was enriched for 1G4 CTL over SIL and is characterized by loss of CD28 and reduced expression of multiple inhibitory receptors (Fig. 8H), suggesting a partial loss of a less active, possibly senescent 1G4 CTL population through spheroid interaction, potentially through antigen exposure in the spheroids.

### Suppressed CTL display inefficient cell couple maintenance and loss of calcium signaling

Cytolysis requires the effective maintenance of CTL couples with target cells (Figs. 5, 6). The inability to maintain a stable cell couple with an activating tumor target cell is a key characteristic of mouse TIL (Ambler et al. 2020). We, therefore, determined the ability of 1G4 SIL to maintain stable cell couples with Mel624 target cells in the presence of 2µg/ml NY-ESO-1 agonist peptide. In comparison to 1G4 CTL, cell couples formed less efficiently (Fig. 9A) but still at a reasonably high frequency of 30±7%. However, first off-interface lamellae were almost instantaneous at 28±1s (Fig. 9B), the majority of cell couples showed translocations (Fig. 9C) and the frequency of detachment was increased compared to 1G4 CTL (Fig. 9D), albeit without a delay in its execution (Fig. 9E). Together these morphological defects resemble CTL activation with a limiting stimulus (Fig. 5B-D, F, G), suggesting that induction of suppression prevents CTL from maintaining stable target cell couples. In contrast to a limiting CTL stimulus (Fig. 5E, S5A), the diameter of the SIL target cell interface however was not reduced (Fig. 9F, S9A). These data suggest that the morphological regulation of reduced SIL function, while sharing many features with CTL activation upon limiting stimulation, is nevertheless distinct, as not further pursued here. The ability of 1G4 SIL to clear F-actin from the interface center was also greatly impaired (Fig. 9G, S9B), while overall interface F-actin accumulation was the same as with 1G4 CTL (Fig. 9H, S9C). Even in the presence of a strong stimulus, 2µg/ml agonist peptide, 1G4 SIL thus were unable to maintain stable cell couples.

**Figure 9.**
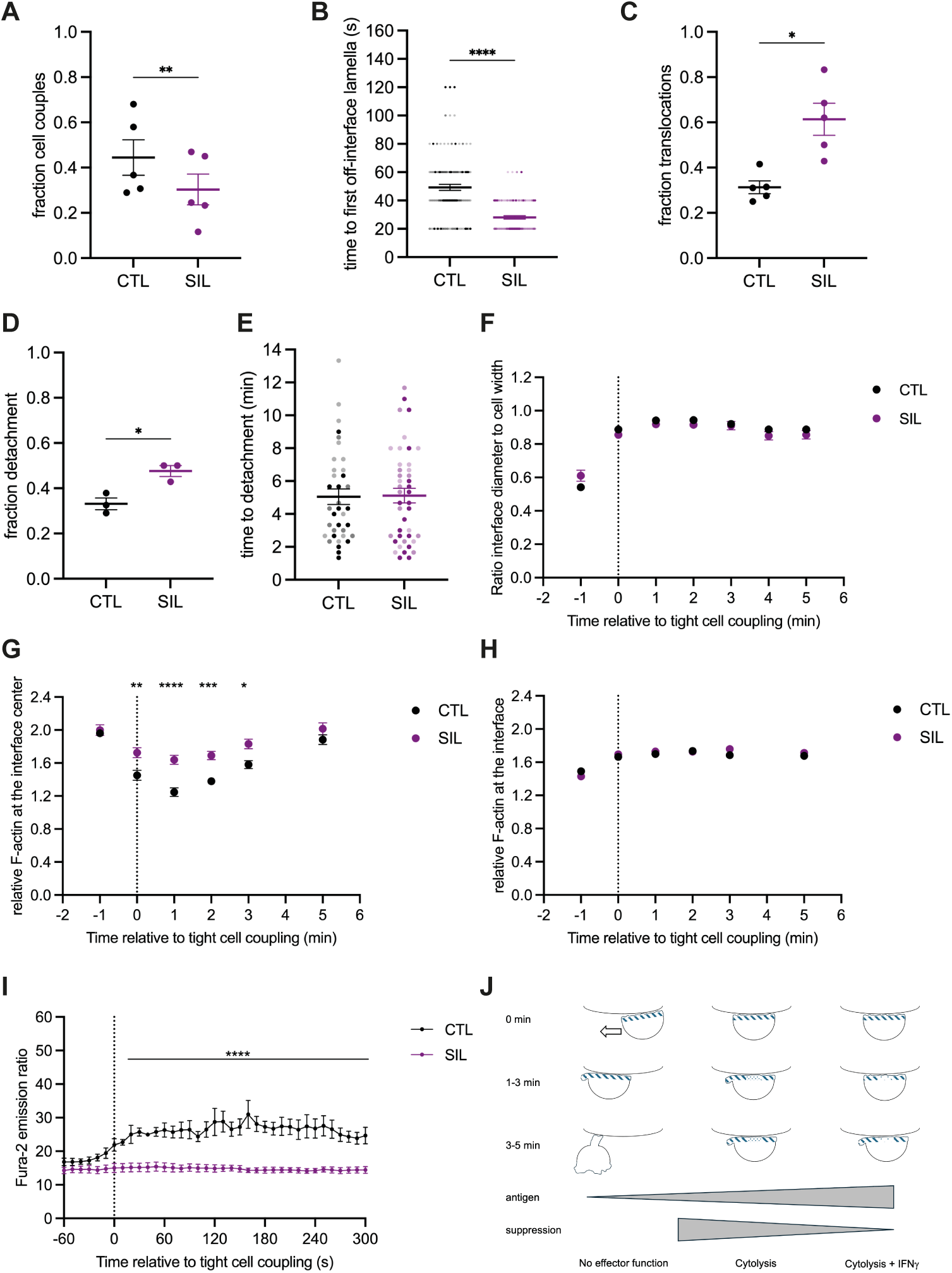
SIL tumor target cell couples are ill maintained **A-F** Characterization of cell morphology in the interaction of 1G4 SIL or CTL with Mel624 cells in the presence of 2µg/ml NY-ESO-1 agonist peptide as mean ± SEM. A Fraction of T cells converting a target cell contact into a tight cell couple. Each symbol is an independent experiment. B Time from tight cell couple formation to first off-interface lamella. Each symbol is a cell couple. Independent experiments are marked by color intensity. C Fraction of CTL with a translocation. Each symbol is an independent experiment. D Fraction of CTL with detachment. Each symbol is an independent experiment. E Time from tight cell couple formation to detachment. Each symbol is a cell couple. Independent experiments are marked by color intensity. F Interface diameters relative to the CTL width. Single cell data in Fig. S9A. 3-5 independent experiments. Statistical significance determined by paired Student’s t-test (A, C, D), Mann-Whitney’s u-test (B) and Two-way ANOVA (F). **G, H** F-actin accumulation at the interface between SIL or CTL expressing the 1G4 TCR with F-tractin-GFP and Mel624 cells in the presence of 2µg/ml NY-ESO-1 agonist peptide relative to F-actin in the entire cell and to the time of tight cell coupling as mean ± SEM. Pooled data from 3 independent experiments. G F- actin accumulation at the central third of the interface. Single cell data in Fig. S9B. Statistical significance determined by Two-way ANOVA. H F-actin accumulation at the entire interface. Single cell data in Fig. S9C. **I** The Fura-2 ratio that is proportional to the cytoplasmic calcium concentration of 1G4 SIL or CTL stimulated with Mel624 cells in the presence of 2µg/ml NY-ESO-1 agonist peptide. Mean ± SEM of 4 independent experiments. Statistical significance determined by paired Two-way ANOVA. Single cell data in Fig. S9D. **J** Scheme of CTL subcellular reorganization in relation to antigen recognition, suppression and effector function. CTL is binding to the target cell membrane above in each individual panel. Time is relative to tight cell couple formation. A large arrow indicates translocation, off-interface lamellae are given at the left of the CTL, if present. F-actin density at the interface and in off-interface lamellae is indicated in dark blue patterning. * p<0.05, ** p<0.01, *** p<0.001, **** p<0.0001.

The elevation of the cytoplasmic calcium concentration is a key proximal T cell signaling step. Even in response to a strong stimulus, such calcium signaling was almost entirely lost in murine TIL and SIL (Ambler et al. 2020). Similarly, when stimulated with Mel624 cells incubated with 2µg/ml NY-ESO-1 agonist peptide, 1G4 SIL did not yield detectable calcium signaling (Fig. 9I, S9D). The highly limited maintenance of 1G4 SIL cell couples together with the loss of calcium signaling are consistent with diminished 1G4 SIL effector function (Fig. 7A, B), similar to murine TIL and support the effective induction of a suppressed phenotype (Fig. 8E).

## Discussion

### Incubation of human CTL with tumor cell spheroids induces a phenotype that closes resembles T cell exhaustion

Incubation of human CTL expressing a transgenic TCR with antigen-presenting tumor cell spheroids induced a loss of CTL function, partially in target cell killing, almost completely in IFNγ secretion (Fig. 7A, B). This could be caused by a reversion of CTL to a resting phenotype or by the induction of exhaustion. When CTL are deprived of IL-2 for 24h in vitro, they retain viability but see a reduction in size, indicative of reversion to a resting phenotype, and drastic changes in their proteome (Rollings et al. 2018). Two of the key changes are loss of expression of CD25 and TIM3 by about 75%. However, CTL incubation with tumor cell spheroids led to retention of high expression levels of CD25 and TIM3 (Fig. 8A-D), arguing against a reversion to a resting phenotype. In contrast, expression of several markers suggested to be defining for exhausted CTL (Winkler et al. 2019) was altered as expected for the induction of CTL exhaustion (Fig. 8E). Comparing marker expression in CTL after spheroid incubation to that before and that in CTL primed for only three days in vitro as fully activated CTL, expression of PD-1, CTLA-4 and LAG- 3 was elevated, expression of CXCR5 and Ki67 was downregulated (Fig. 8E). While the fraction of CTL expressing TIGIT wasn’t increased (Fig. 8A), the MFI was doubled in comparison to CTL primed for only three days (Fig. 8B). All these changes are consistent with induction of CTL exhaustion upon spheroid co-culture. The only difference from the expected results was that TCF-1 expression was slightly elevated rather than decreased, albeit at the low level of about 10% of CTL expressing TCF-1 after spheroid co-culture (Fig. 8E). As this is the frequency of TCF-1^+^ CTL seen in patients (Sade-Feldman et al. 2018, Jansen et al. 2019, Beltra et al. 2020), the increase in TCF-1 expression upon spheroid co-culture likely indicates low expression of TCF-1 in the CTL to be incubated with the spheroids rather than an unusual induction of TCF-1 expression.

Comparing marker expression in the spheroid suppressed CTL to that in tumor-infiltrating CD8^+^ T cells from melanoma patients, high expression of PD-1 is shared (Fig. 8A, D, E). However, alignment with other markers such as CTLA-4 and TIM-3 is weaker. A possible explanation is that the TIL are substantially less activated as indicated by lower CD25 expression (Fig. 8A, C). While the in vitro CTL are homogeneously antigen-reactive and have interacted with antigen in the spheroids, a substantial fraction of CD8^+^ T cells in tumors are bystander cells (Simoni et al. 2018, Meier et al. 2022), likely leading to more variable activation phenotypes. With respect to the time needed to induce CTL exhaustion, exhausted CTL reside in tumors for days (Li et al. 2022). However, hallmarks of CTL dysfunction can be established in hours (Rudloff et al. 2023), as used in our spheroid experiments. In addition, the in vitro CTL cultured for seven days in the presence of anti-CD3/CD28 beads used for the spheroid co-culture already showed some upregulation of exhaustion markers such as PD-1, CTLA-4 and LAG-3 as well as downregulation of Ki67 (Fig. 8E). Some of these changes were dependent on the continued presence of the anti-CD3/CD28 beads (Fig. S8E). It thus is conceivable that the effective induction of a CTL phenotype closely resembling exhaustion by spheroid co-culture is made more readily achievable by the partial polarization of the in vitro CTL towards exhaustion before spheroid incubation. This suggestion needs to be tested in the future.

Overall, our characterization of CTL after spheroid interaction with respect to function and marker expression has established close similarity with tumor-exhausted CTL. This similarity is further corroborated by an inability of the human SIL to effectively maintain stable cell couples and a complete loss of calcium signaling (Fig. 9), a phenotype initially described in murine TIL (Ambler et al. 2020). While we can’t map the in vitro SIL in detail onto the in vivo progression from precursor to terminally exhausted TIL, the retention of some cytolytic functionality suggests that the SIL most closely resemble the intermediate bulk stage of cytolytic exhausted TIL. Thus, the spheroid-suppressed CTL thus should be a valuable in vitro model for the investigation of mechanisms of CTL exhaustion and therapeutic means to overcome it, as already initiated (Alamir et al. 2023).

### Spheroid properties in the induction of CTL suppression

The ability of spheroids to effectively induce CTL suppression upon co-culture raises the question which spheroid properties are key to this ability. We have not systematically investigated this question here, but can offer suggestions: mechanical properties, altered metabolism and sustained antigen presentation. Tumor cell biology changes upon growth in 3D structures. In CRISP screens to identify cancer driver genes, hits identified in 3D spheroids matched in vivo biology more closely than hits identified in 2D tumor cell culture (Han et al. 2020). A key element of altered tumor cell biology is increased stiffness in a 3D environment (Broders-Bondon et al. 2018). As three-dimensional structures spheroids force CTL to operate in a constrained environment. In vivo, biomechanical stress as sensed by CTL using Piezo1 enhances the induction of T cell exhaustion (Zhang et al. 2024). The more constrained biomechanical environment of spheroids thus is a likely contributor to the induction of CTL suppression.

In murine renal carcinoma cells, growth in 3D spheroids alters metabolic properties of the tumor cells. Expression of multiple amino acid transporters on the tumor cell surface is downregulated upon tumor cell growth in spheroids while expression of the glucose transporter Slc2a1 is upregulated (ref), suggesting more efficient glucose uptake. Glucose limitation in a 3D microenvironment can inhibit CTL effector function (Chang et al. 2015, Lim et al. 2020). As an additional element of metabolic regulation, PD-1 engagement leads to impaired mitochondrial function as a key contributor to T cell exhaustion (Lim et al. 2020, Vardhana et al. 2020, Yu et al. 2020). We have shown that PD-1 is further upregulated upon CTL co-culture with spheroids (Fig. 8A, E). Additional tumor metabolites that may be similarly induced or limiting in spheroids, such as D-2-hydroxyglutarate or taurine, respectively, could also promote T cell exhaustion (Notarangelo et al. 2022, Cao et al. 2024). Altered tumor cell metabolism could act directly on the CTL through metabolic competition and regulation or indirectly, as it is linked to changed biomechanics (Park et al. 2020). Altered CTL metabolism as forced by changes in tumor cell metabolism and inhibitory receptor expression induced in spheroids thus is also likely to contribute to the induction of CTL suppression.

Persistent antigen exposure is a well-established inducer of CD8^+^ T cell exhaustion (Barber et al. 2006). However, this may not require tumor cell growth as spheroids. Here we have shown that persistent interaction of CTL with anti-CD3/CD28 beads can induce a partially suppressed CTL phenotype (Fig. S8C-E). Small amounts of endogenously processed antigen in spheroids were as effective in inducing CTL suppression as a high concentration of exogenous agonist peptide (Fig. 7C, D), arguing that antigen amounts are not critical for the induction of suppression. Together these data suggest that persistence of antigen presentation in spheroids alone is unlikely to be the key driver of the induction of CTL suppression. In summary, we suggest a combinatorial model of the induction of CTL suppression in spheroids were altered biomechanics and metabolism in conjunction with persistent antigen presentation and potential additional factors that we have not considered here together drive efficient induction of CTL suppression, as to be verified experimentally in the future.

### CTL effector function correlates more with the effective maintenance of target cell couples than with their formation

The execution of CTL effector function requires a series of well-characterized changes in CTL subcellular organization. An initial contact with a potential target cell as mediated by individual lamellae needs to be converted into a tight interface almost as wide as the CTL as supported by CTL F-actin accumulation at the interface. The MTOC needs to reorient from behind the nucleus to the cellular interface, bringing along the secretory machinery (Kupfer et al. 1989, Frazer et al. 2021). The density of F-actin at the center of the cellular interface needs to be reduced to allow the effective release of lytic granules at the interface center (Ritter et al. 2015). CTL can execute their effector function without the formation of supramolecular signaling structures at the interface with target cells (Purbhoo et al. 2004, Purtic et al. 2005). Supramolecular complex formation was therefore not assessed here. Subsequent resolution of the cell couple is important to allow sequential killing of multiple target cells (Halle et al. 2016, Weigelin et al. 2021). While this series of subcellular rearrangements is well defined for cytolytic killing, it is less well understood how these events relate to cytokine secretion. Here we have investigated CTL subcellular polarization in relation to cognate antigen recognition and CTL effector function. The formation of a wide, F-actin-supported interface with a target cell in a sizable fraction of CTL did not require cognate antigen (Fig. 5B, S5A, 6C). While cell coupling was more efficient in the presence of antigen, about 20% of initial target cell contacts were converted to wide interfaces even in the absence of antigen. At best minimal CTL effector function was associated with such cell coupling (Fig. 3B, C). Therefore, at least part of the decision whether to trigger effector function must occur in the progression through the subsequent subcellular reorganization steps. Indicators of efficient CTL reorganization are the late, more than a minute, first formation of destabilizing lamellae that point away from the cellular interface with the target cell, the lack of translocation of the CTL over the target cell surface away from the initial site of binding, a wide target cell interface and a substantial reduction in F-actin content at the interface center. All four required a higher amount of agonist and needed to occur for a CTL to commit to IFNγ secretion (Fig. 6F). Cytolysis was more permissive. Here rapid occurrence of the first off-interface lamellae, intermediate interface width and only partial F-actin clearance at the interface center still allowed for substantial target cell killing. However, an almost complete absence of translocations was still required (Fig. 6F). For such cytolysis, small amounts of endogenously processed antigen sufficed. Together, these data suggest a model of a stepwise generation of CTL effector function in response to antigen (Fig. 9J). A base level of tight cell coupling as supported by F-actin accumulation occurs even in the absence of cognate antigen and is not associated with effector function. If the CTL can remain at the site of initial cell coupling for a few minutes, even with only partial subcellular reorganization, as enabled by limiting amounts of endogenously processed antigen, cytolysis is triggered. Killing by CTL has been shown to occur on the time scale of not more than one minute (Regoes et al. 2007). Only maintenance of a fully reorganized CTL with substantial F-actin clearance at the interface center, late off-interface lamella, absence of translocations and detachment only after several minutes requiring a larger amount of antigen allows for IFNγ secretion. A less stringent antigen threshold for cytolysis versus IFNγ secretion has been described before (Betts et al. 2004). While we have focused here on antigen amounts, the role of antigen affinity on CTL subcellular organization has been extensively characterized with key roles of cell couple duration, MTOC and granule reorientation and F-actin clearing at the interface center (Frazer et al. 2021). Together with our data here, these data are consistent with the notion that antigen affinity and amount jointly control T cell responses (Pettmann et al. 2021).

In the context of this model, the induction of CTL suppression by spheroid co-culture prevents the establishment of a fully reorganized CTL even in response to high concentrations of agonist (Fig. 9). Suppressed CTL were characterized by rapid off-interface lamellae, an increased frequency of translocation and detachment as well as impaired clearance of F-actin at the interface center. According to our model, such defective subcellular reorganization is incompatible with IFNγ secretion and allows only reduced cytolysis, as confirmed experimentally (Fig. 7A, B). Progressive subcellular reorganization thus offers a unified model of the regulation of CTL function where increasing antigen-mediated stimulation gradually drives more effective CTL reorganization and function while the induction of suppression counteracts it.

## Materials and Methods

### Human blood samples

Blood buffy coats from anonymous donors were purchased from NHS-BT with human work approved by the London-Riverside Research Ethics Committee under reference number 20/PR/0763.

### Antibodies

Antibodies and staining reagents used are described in the order: antigen, fluorescent label, clone, supplier, dilution/concentration, RRID:

Panel for flow cytometry

Human CD25, BV421BD, M-A251, BD Bioscience, 1:100, RRID:AB_11154578 Human CD25, BV421BD, 2A3, BD Bioscience, 1:100, RRID:AB_2738555

Human CD8α, Pacific Blue, RPA-T8, BioLegend, 1:25, RRID:AB_493111 Human CD28, BV480, CD28.2, BD Bioscience, 1:100, RRID:AB_2739512

Human CD4, Starbright V570, RPA-T4, Bio-Rad, 1:50, RRID:AB_3099780

Human CD3ε, Super Bright 645, OKT3, eBioscience, 1:20, RRID:AB_2662368 Human CD185 (CXCR5), BV785, J252D4, BioLegend, 1:50, RRID:AB_2629527

Human CD366 (TIM-3), Super Bright 600, F38-2E2, eBioscience, 1:100, RRID:AB_26882087 Human CD45RA, R718, HI100, BD Bioscience, 1:100, RRID:AB_2916420

Human CD279 (PD-1), APC-Cy7, EH12.2H7, BioLegend, 1:100, RRID:AB_10900982

Human CD45, NovaBlue 610-30S, 2D1, eBioscience, 1:100, RRID:AB_3098021 Human CD352 (SLAMF6/Ly108), biotin, REA339, Miltenyi, 1:30, RRID:AB_2657676

Streptavidin, Star Bright Blue 675, Bio-Rad, 1:85

Human CD272 (BTLA), BB700, J168-540, BD Bioscience, 1:50, RRID:AB_2743523 Human CD152 (CTLA-4), PE-Fire 640, BNI3, BioLegend 1:20, RRID:AB_2924561 Human TIGIT, PE-vio770, REA1004, Miltenyi, 1:100, RRID:AB_2751339

Human CD223 (LAG-3), PE-Fire 810, 7H2C65, BioLegend, 1:100, RRID:AB_2927867 Human CD14, RB545, M5E2, BD Bioscience, 1:100, RRID:AB_2691229

Human CD19, RB545, SJ25C1, BD Bioscience, 1:100, RRID:AB_2688586 Human Ki-67, BV711, Ki-67, BioLegend, 1:100, RRID:AB_11218996

Human Foxp3, PE, 259D, BioLegend, 1:50, RRID:AB_492983 Human Foxp3, PE, PCH101, eBioscience, 1:50, RRID:AB_1518782

Human TCF1, Alexa Fluor 647, 7F11A10, BioLegend, 1:50, RRID:AB_2566619 Human T-bet, RB780, O4-46, BD Bioscience, 1:100, RRID:AB_2688938

Live Dead Red, Invitrogen, 1:20,000

Individual antibodies and staining reagents for flow cytometry

Human NY-ESO-1, PE, D1Q2U, Cell Signaling Technology, 1:50, RRID:AB_2799691 Human TCR Vβ20, PE, ELL1.4, Beckman Coulter, 1:30, RRID:AB_131328

Human TCR Vβ13.1, PE, IMMU222, Beckman Coulter, 1:30, RRID:AB_131326

Tetramer HLA-A*0201, MART-126-35 peptide with the A27L mutation, PE, gift from L. Wooldridge (U. Bristol), 10µg/ml

For Western Blotting

Human NY-ESO-1, E978, Santa Cruz Biotechnology, 1:200, RRID:AB_784921 Human MART-1, A103, Santa Cruz Biotechnology, 1:200, RRID:AB_627912

### Human Cell culture

Human A375 melanoma (RRID:CVCL_0132), Mel624 melanoma (RRID:CVCL_8054) and NCI- H1755 non-small cell lung carcinoma (RRID:CVCL_1492) cells were stably transfected to express mCherry or tdTomato (Shaner et al. 2004) using a plasmid derived from pIRES Hygro (Addgene). A375 and Mel624 cells were maintained in high glucose DMEM with 10% FBS, 2mM Glutamine, 1mM pyruvate (DMEM complete medium). NCI-H1755 were maintained in RPMI1640 with 10% FBS and 2mM Glutamine (RPMI complete medium). Medium for the growth of mCherry or tdTomato transfectants was supplemented with 250μg/ml Hygromycin.

Blood buffy coats were obtained from healthy donors. PBMC were isolated by density gradient centrifugation using Ficoll-Paque^TM^ (Sigma-Aldrich). These cells constitute the flow cytometry ‘PBMC, unstimulated’ sample. For the ‘PBMC, stimulated sample’ these cells were incubated with anti-CD3/CD28 beads as described below for 3d only. PMBC were cryopreserved at a concentration of 2.5x10^7^ cells/ml. To isolate CD8^+^ T cells, buffy coat cryopreserved vials were thawed, the cells were washed twice with RPMI 1640 with 10% FBS, 2mM L-glutamine, 50µM β-mercaptoethanol and resuspended in ice cold MACS buffer. CD8^+^ T cells were purified by magnetic enrichment for CD8^+^ cells using CD8 MicroBeads (Miltenyi Biotech). CD8^+^ T cells were activated using CD3/CD28 Dynabeads (Life Technologies) at bead-to-cell ratio of 1:1 in human IL-2 medium (X-VIVO 15, serum-free hematopoietic cell medium, with 2mM L-Glutamine and gentamicin (Lonza) supplemented with 5% Human AB serum (Valley Medical), 10mM neutralized N-acetyl L-Cysteine (Sigma-Aldrich), 50μM β-Mercaptoethanol (Gibco, Thermo Fisher), and 30U/ml rh-IL-2 (NIH/NCI BRB Preclinical Repository – human IL-2 medium) and incubated overnight at 37 °C and 6% CO2.

The 1G4 and MEL5 TCRs, as stabilized with an additional disulfide bridge in the constant domains (Cohen et al. 2007), were expressed in primary human T cells using a pHR_SFFV - based lentiviral vector (RRID_Addgene79121) with an expression cassette of alpha chain-P2A- beta chain-P2A-GFP (as a sorting marker) or F-tractin-GFP (for F-actin imaging). For the generation of lentiviral particles, HEK 293T cells (RRID:CVCL_0063; Lenti-X 293T cells, Takara) were maintained in DMEM complete medium. 1.5×10^6^ Lenti-X 293T cells were seeded in 5ml DMEM complete medium in 60x15 mm Primaria culture plates (Corning) 24h before transfection. Cells were transfected with a total of 4.5 µg plasmid DNA using Fugene HD (Promega): 0.25µg envelope vector pMD2.G (RRID:Addgene_12259), 2µg of packaging plasmid pCMV-dR8.91 (Creative Biogene), and 2.25µg of the pHR_SFFV-based transfer vector. 48h after transfection, virus containing medium was collected and filtered through a 0.45µm nylon filter. The MEL5 TCR as stabilized by swapping the human constant domains against murine ones (Cohen et al. 2006) was expressed using lentiviral constructs and procedures as described in (Kalaitsidou et al. 2023) with an expression cassette of alpha chain-P2A-beta chain-T2A-FRα CoStAR mCherry-E2A-F-tractin-GFP. For lentiviral infection of CTL, after 24h of setting up the primary human T cell culture, 1x10^6^ T cells were mixed with 500-700µl lentivirus-containing medium in a 24-well plate medium in presence of 8µg/ml Polybrene (Sigma-Aldrich) and centrifuged for 1.5h at 2500rpm, 37 °C. After spinduction primary CD8^+^ T cells were resuspended in human IL-2 medium. Cells were maintained at density of less than 2x10^6^ cells/ml and if necessary spilt back to a density of 0.5-1x10^6^ cells/ml. For some experiments as indicated, CD3/CD28 Dynabeads were removed before the spinduction or two days thereafter.

Isolation of tumor-infiltrating lymphocytes. Around 0.5g of freshly dissected specimen was collected from melanoma patients undergoing surgery at the UHBW Hospital. Specimen were maintained in 20ml of MACS® Tissue Storage Solution (Miltenyi Biotec) protected from light at 2−8°C. Samples were processed within 4 hours of collection. Tumour samples were dissected into 2 to 4mm pieces and enzymatically digested using the Human Tumour Dissociation Kit (Miltenyi Biotec). Samples were filtered to single-cell suspension by passing through a 70µm cell strainer, centrifuged at 1500rpm for 5 minutes and resuspended in PBS to 1x10^6^ cells/ml.

### Spheroids and SILs

A375 tdTomato or Mel624 tdTomato cells were resuspended at a concentration of 1×10^5^ cells/ml, mixed with Matrigel (Corning) at 4°C, seeded in a 24-well plate at a final concentration of 500 cells per Matrigel dome, and left to solidify for 10min at 37°C. 2ml DMEM complete medium was added to each well and cells incubated at 37°C for 11 days as the default and up to 17 days in the characterization of spheroid properties. To determine spheroid properties, bright field images were acquired on indicated days of spheroid culture and the area of the spheroid cross-section, spheroid circularity and roundness were measured using the particle analysis function of ImageJ/Fiji. NCI-H1755 cells were prepared in RPMI complete medium supplemented with 2.5 % Matrigel. The cells were seeded in a 96-well round-bottom plate pre-coated with 1% agarose at a concentration of 2000 cells per well. The plate was then centrifuged at 1000 rpm for 5 mins to encourage cells to cluster. Cells were maintained in culture for 5-8 days.

For the generation of spheroid-infiltrating lymphocytes (SIL), each Matrigel dome was washed twice in PBS and incubated for 1h with 1ml of Cell Recovery Solution (Corning). Spheroids were collected in a 15-ml Falcon tube and pulsed with NY-ESO-1 peptide at a final concentration of 2μg/ml for 1h or left unpulsed. Spheroids were re-embedded in Matrigel together with 5×10^6^ 1G4 CTL per Matrigel dome. Matrigel domes were dissolved for analysis of spheroid-infiltrating T cells after 16h: Spheroids were washed twice in PBS and incubated with 1ml of Cell Recovery Solution (Corning). Spheroids were collected, washed through a 40μm sieve and then disaggregated to retrieve T cells in 500μl of imaging buffer for immediate FACS sorting.

To determine CTL infiltration into spheroids by live cell imaging, spheroids were dissociated from Matrigel and resuspended into fresh Matrigel at a concentration of ∼8 spheroids/µl. 50µl of the spheroid-Matrigel suspension was separated into Eppendorf tubes, followed by the addition of 500,000 human sorted CTL per tube. 50µl of Matrigel, containing spheroids and T cells, was plated into each well of a 24-well tissue culture plate. After Matrigel had set, 1ml of Fluorobrite medium (ThermoFisher) was added to each well, containing 1.5µM DRAQ7 viability. Images were acquired every 2h post-plating CTL with spheroids in 3µm z steps from the bottom of the spheroid to its widest point, usually 40 steps, for 12h using a Leica SP8 AOBS confocal microscope with a 10x HC PL Fluotar lens (NA=0.3). Spheroids and SIL were segmented using a custom Fiji Image J script as described before (Edmunds et al. 2022). Distance of each T cell from the spheroid surface was automatically calculated using the script.

### Cytolysis and IFNγ secretion

For imaging-based cytotoxicity assays, the IncuCyte™ Live Cell analysis system and IncuCyte™ ZOOM software (Essen Bioscience) were used to quantify target cell death. 1x10^6^ Mel624, A375 or NCI-H1755 cells transfected to express the fluorescent protein mCherry or tdTomato were either untreated or pulsed for 1h with the indicated concentration of HLA-A*02:01 NY-ESO-1157- 165 (SLLMWITQC), HLA-A*02:01 MART-126-35 with the A27L mutation (‘ELA’)(ELAGIGILTV) or HLA-A*02:01 MART-126-35 with the E26F, A28T double mutation (‘FAT’)(FATGIGILTV). Cells were suspended in 5ml Fluorobrite medium (ThermoFisher) with 10% FBS, 2mM L-glutamine, 50µM 2-mercaptoethanol to a concentration of 15,000 cells/50μl. Cells were plated in each well of a 384 well Perkin-Elmer plastic-bottomed plate and incubated for 4h to adhere. 10,000 - 40,000 CTL that had been FACS sorted for indicated expression of GFP were added per well to the plate in 50μl Fluorobrite medium, yielding 1:1 to 4:1 effector to target ratios, respectively.

Images were taken every 15min for 14h at 1600ms exposure using a 10x (NA=0.3) lens. The total red object (mCherry/tdTomato target cell) area (µm^2^/well) was quantified at each time point. A change in red area was determined as the linear gradient of the red object data at its steepest part for 6h. The CTL killing rate was calculated as the difference in such change in red area in the presence and absence (to account for tumor cell proliferation) of CTL in the same 6h time window.

To determine IFNγ secretion, supernatants at the end of the imaging-based cytolysis assays were collected and frozen. The human IFNγ OptEIA ELISA Kits (BD Biosciences) was used according to manufacturer’s instructions. Briefly, wells of a 96-well Maxisorb Nunc-Immuno plate (ThermoFisher) were coated with 100µl anti-human IFNγ monoclonal antibody diluted in 0.1M Na2CO3 coating buffer at a 1:250 dilution and incubated overnight at 4°C. Plates were blocked using PBS with 10% FBS for 1h. Samples were diluted in ELISA dilution buffer at 1:20 and incubated for 1h at room temperature. 100µl of working detectors (Biotinylated anti-human IFNγ monoclonal antibody + Streptavidin-horseradish peroxidase conjugate) were incubated for 1h at RT before detection. Absorbance at 450nm and 570nm was measured within 20min. Samples were measured in duplicate or triplicate.

### Imaging of CTL and SIL

Prior to imaging, CTL and SIL were resuspended in ‘imaging buffer’ (10% FBS in PBS with 1mM CaCl2 and 0.5mM MgCl2). As target cells, 1x10^6^ Mel624 or A375 melanoma cells were pulsed with the indicated peptide (as listed above) at a final concentration of 2µg/mL for 1 hour or left unpulsed. Glass bottomed, 384-well optical imaging plates (Brooks life science systems) were used for all imaging experiments. Imaging of F-actin distributions and CTL morphology was done at 37°C using an Olympus IXplore SpinSR confocal system incorporating a Yokogawa CSU-W1 SoRa spinning disk. A 60x oil-immersion lens (NA=1.5) was used for all imaging experiments, unless otherwise stated. Images were acquired for 15 minutes. Every 20 seconds, a z-stack of 53 GFP images (0.25µm z-spacing) was acquired, as well as a single, mid-plane differential interference contrast (DIC) reference image.

For imaging the elevation of the cytoplasmic calcium concentration, 1G4 T cells were incubated with 2 µM Fura-2 AM (Molecular Probes) for 30min at room temperature in imaging buffer. 1G4 CTL were washed twice thereafter. Because of limiting cell numbers 1G4 SIL were only washed once. 1G4 T cells were activated with Mel624 APCs as described above and three images were acquired every 10s for 15min, one bright field image, one fluorescence image with excitation at 340nm and one fluorescence image with excitation at 380nm. Imaging data were acquired at 37°C using a 40x oil objective (NA=1.25) on a Leica DM IRBE-based wide filed system equipped with Sutter DG5 illumination and a Photometrics Coolsnap HQ2 camera.

### Analysis of live cell imaging data

Using Fiji/ImageJ (Schindelin et al. 2012, Rueden et al. 2017) for analysis of DIC images, tight cell couple formation was defined as the first time point at which a maximally spread immune synapse formed, or 40s after initial cell contact, whichever occurred first. To assess CTL and SIL morphology in cell couples with tumor target cells, every DIC frame after tight cell couple formation was assessed for the presence of off-synapse lamellae, defined as transient membrane protrusions pointing away from the immune synapse, followed by retraction. To determine CTL translocation over the tumor cell surface, the position of the immune synapse on the tumor target cell was compared to the position at cell coupling. If the T cell had migrated by a distance greater than the diameter of the immune synapse, this was classed as translocation. The formation of a uropod that is bound to the tumor cell at the interface together with lamellae at the CTL side opposite of the interface was classed as ‘detachment’. The diameter of the CTL target cell interface was measured in the DIC image as a straight line from one end of the interface to the other. It was normalized by division through the CTL diameter, measured as a straight line across the widest part of the CTL parallel to the interface.

For analysis of F-actin distributions as imaged with F-tractin-GFP, enrichment at the interface and interface center were measured relative to F-actin across the entire cell using Fiji/ImageJ. F-tractin-GFP enrichment across the interface relative to the entire cell was determined in the maximum intensity z-projection of the GFP z-stack. The interface was defined as the 10% of the area of the T cell closest to the T cell/target cell interface. The average fluorescence intensity of the interface area and the entire cell were measured at four to six timepoints after subtracting the mean of ten background fluorescence readings. F-tractin-GFP enrichment at the centre of the interface relative to the entire cell was determined in the midplane of the GFP z-stack of the

T cell of interest. The area of the interface centre was defined as the middle third of the interface as defined above, that is as 3.5% of the total cell midplane area. The average fluorescence intensity of the interface center and the entire cell were measured at four to six timepoints after subtracting the mean of ten background fluorescence readings.

For analysis of the Fura-2 imaging experiments using Fiji/ImageJ, rolling ball background fluorescence was subtracted from the fluorescence data and the ratio of the Fura-2 images upon excitation at 340nm versus 380nm was calculated and multiplied by 100 to fit into the 8-bit display scale. Average ratio within a circular region of interest of the dimensions of the T cell was determined over time for each T cell.

### Flow cytometry

21-color panel: 1x10^6^ cells were first stained with a 1:10,000 dilution of the LIVE/DEAD™ Fixable Red Dead Cell Stain (ThermoFisher) for 15-30 minutes in the dark at room temperature (RT) and washed with FACS buffer (PBS, 0.5% BSA, 2.5mM EDTA). Fc receptors were blocked with 10µl of a 1:20 dilution of Fc Blocker (ThermoFisher) in the dark at RT for 10 minutes. 40µl of biotinylated antibody was added and incubated at 4°C for 30 minutes. The sample was washed with FACS buffer and incubated with 10µl of a 1:20 dilution of Monocyte Blocker (BioLegend) in the dark at RT for 10 minutes. 40µl of an antibody master mix against cell surface proteins as listed in the antibodies section was added for the total volume of 50µl, incubated at 4°C for 30 minutes and washed with FACS buffer. Cells were fixed and permeabilized using the True-Nuclear™ Transcription Factor Buffer Set (BioLegend). Cells were stained for intracellular proteins with an antibody master mix as listed in the antibodies section for 30 minutes at 4°C. Cells were washes with permeabilization buffer and resuspended in FACS buffer and kept at 4°C until the next day. For the flow cytometry data acquisition, samples were run on four-laser Cytek® Aurora system (4L V/B/YG/R) on low to medium flow rates (15 - 30 µl/min) and analysed using SpectroFlo® software with autofluorescence extraction and live unmix during sample acquisition.

The data for the unmixed samples were processed using the FlowJo software (v.10.10.0). Positive gates were set using fluorescence-minus-one data.

The same protocol, live/dead stain, Fc block, cell surface antibodies, permeabilization, internal antibodies, was used for single antibody flow cytometry experiments with steps omitted as allowed by the antigen to be stained. For tetramer staining, some samples were treated with 50nM of the tyrosine kinase inhibitor dasatinib (a gift from L. Wooldridge, U. Bristol) to enhance tetramer staining (Lissina et al. 2009).

For cluster analysis, samples were gated for live, singlet, CD45^+^CD3^+^CD19^−^CD14^−^CD8^+^ cells. The FlowJo 10.10.0 clean plugin was used to reduce technical noise. Files for experimental repeats were concatenated with the FlowJo concatenation tool. CD8^+^ cells for each experimental repeat were down sampled to 18,500 cell and analyzed for CD25, CD28, CD185, CD366, CD45RA, CD279, CD352, CD272, CD152, TIGIT, CD223, Ki-67, FOXP3, TCF1, T-BET.

t-SNE (t-Distribution Stochastic Neighbor Embedding) was employed to map the high-dimensional cellular data into a two-dimensional space using: iteration: 1000, perplexity: 30, K-nearest neighbors’ algorithm: Exact (vantage point tree), and gradient algorithm: Barnes-Hut. Cells were clustered by implementing FlowSOM v. 3.0.18 with 10 metaclusters and 10 × 10 grid size, all other parameters were left at the default settings.

### Data Availability

Raw data are available upon request.

## Acknowledgements

We acknowledge support from the University of Bristol Flow Cytometry and Wolfson BioImaging core facilities. We thank Dr. Helen Winter (Bristol Cancer Institute) for coordinating access to melanoma patient samples and Prof. Linda Wooldridge (U. Bristol) for the MART-1/HLA-A*0201 tetramer.

## Additional information Funding sources

This work was supported by grants from the MRC (MR/W006308/1 to TG for the GW4 BIOMED MRC DTP), InstilBio (to JSB and CW) and Immunocore (to CJH and CW). HA, AmA, AbA and MA were supported by the Ministry of Education of Saudi Arabia.

## Author contributions

AmA, HA, LH Investigation, Formal Analysis, Methodology, Writing Review & Editing; TG, AbA, PHC, MA, YS, MA, LRM, Investigation, Formal Analysis; AH, Methodology, JSB, CJH, Funding Acquisition, Writing Review & Editing, CW Investigation, Formal Analysis, Visualization, Funding Acquisition, Writing Initial Draft, Writing Review & Editing

