## Supplementary figures for "Interaction with tumor cell spheroids induces suppression in primary human cytotoxic T cells"

**Figure S1**

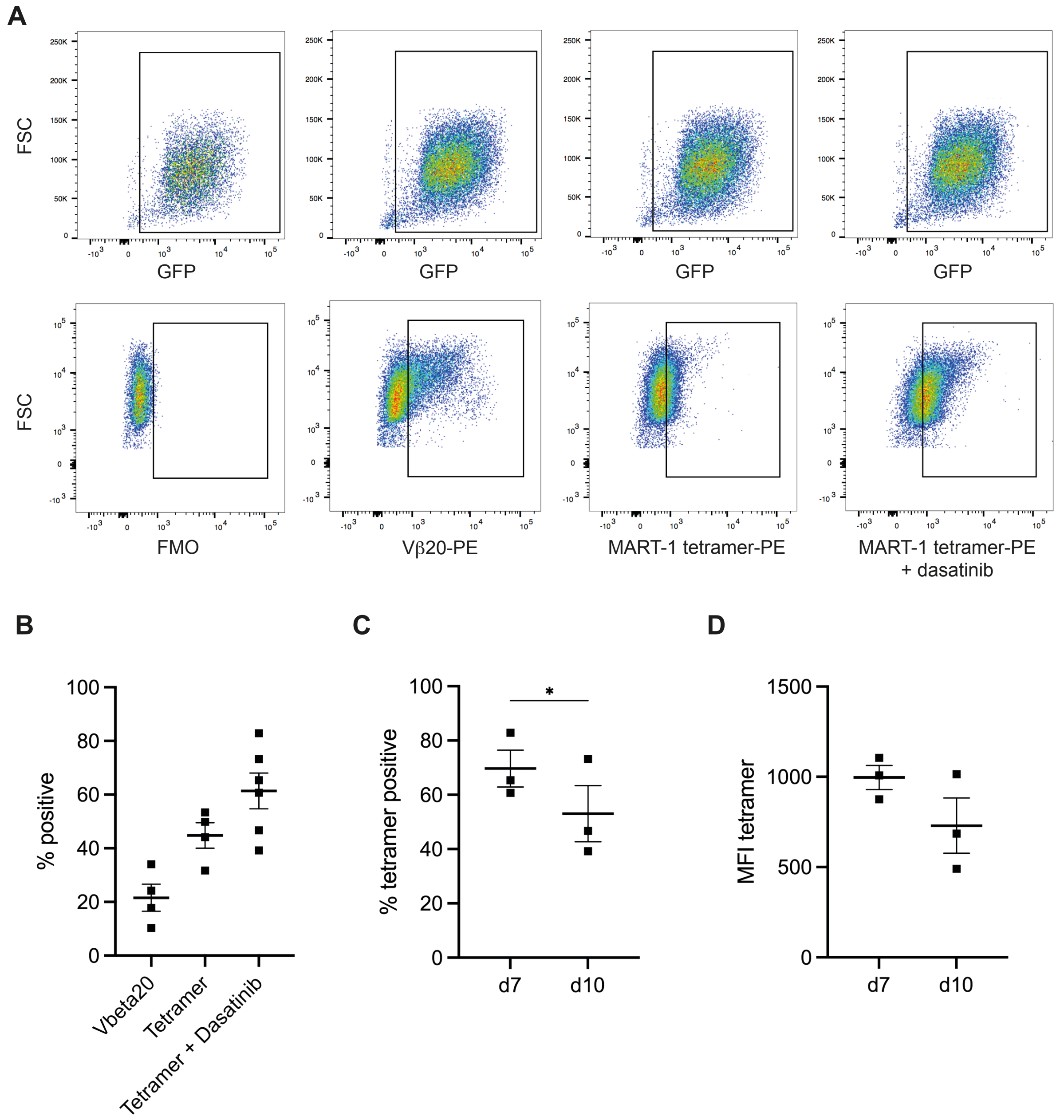


**Transient expression of the MEL5 TCR in human primary CTL**

**A** Representative staining data of primary human CTL transduced to express the MEL5 TCR. The top row shows GFP expression as the transduction marker. The bottom row shows staining for the MEL5 TCR as indicated below each panel. Shown are all live cells. 1 of ≥4 independent experiments. **B** Percent live CTL transduced to express the MEL5 TCR on day 7 in the $GFP^{+++}$ gate that are positive for Vβ20, the MART-1/HLA-A*0201 tetramer or the tetramer in the presence of dasatinib as mean ± SEM. 4 to 6 independent experiments. **C, D** Percent live CTL transduced to express the MEL5 TCR on day 7 and 10 in the $GFP^{+++}$ gate that are positive for the MART-1/HLA-A*0201 tetramer in the presence of dasatinib and MFI of tetramer stain. 3 independent experiments. Statistical significance determined by paired Student's t-test. C Percent live cells, D MFI. * $p<0.05$.

**Figure S3**

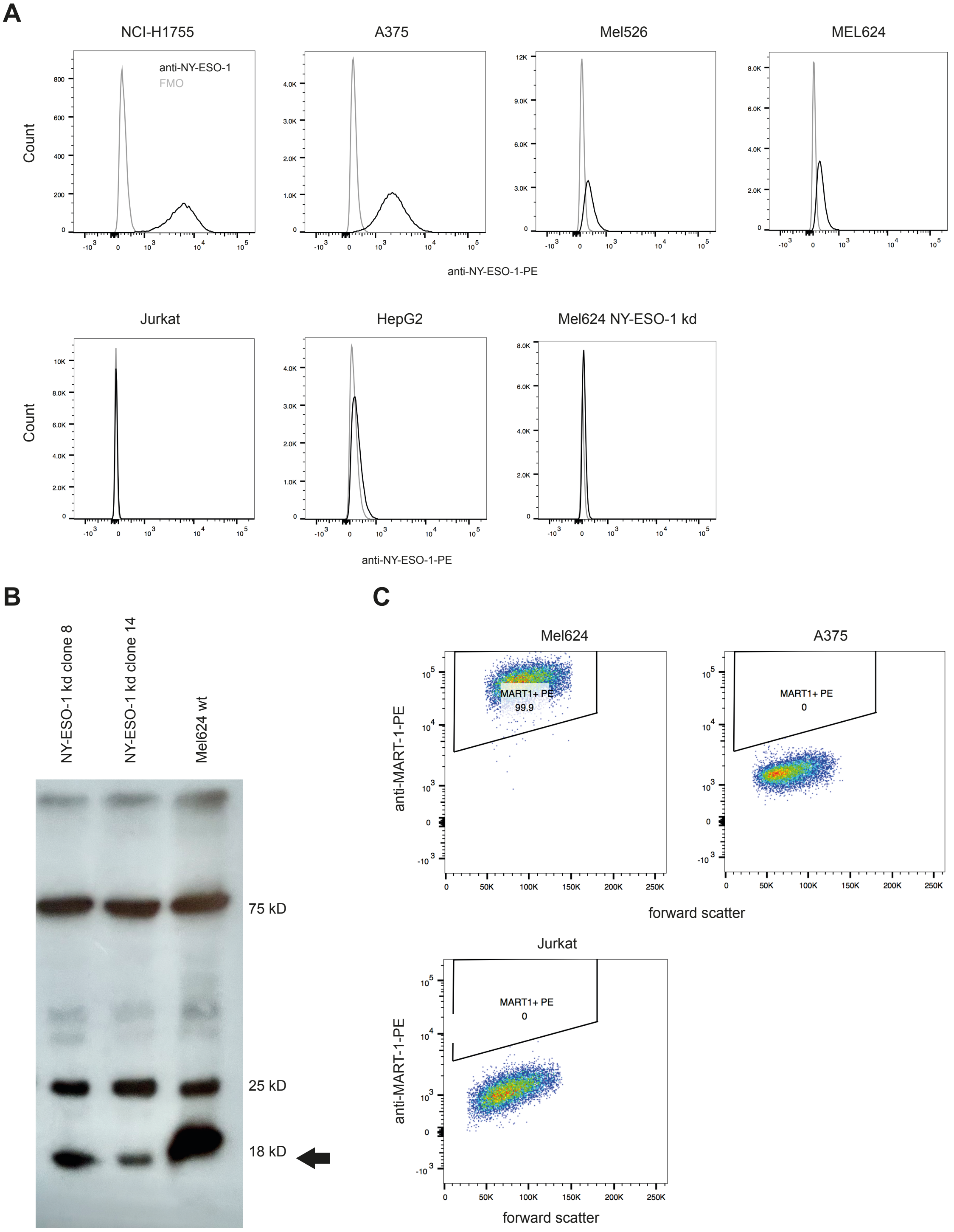


**Expression of NY-ESO-1 and MART-1 in tumor cell lines**

**A** Representative staining of the indicated cells for NY-ESO-1 (black lines) in comparison to fluorescence minus one (FMO)(grey lines). 1 representative experiment of up to 4. **B** Representative Western Blot of Mel624 NY-ESO-1 knock down clones. 1 of 2 independent experiments. **C** Representative staining of the indicated cell lines for MART-1. 1 of 2 independent experiments.

### Figure S4

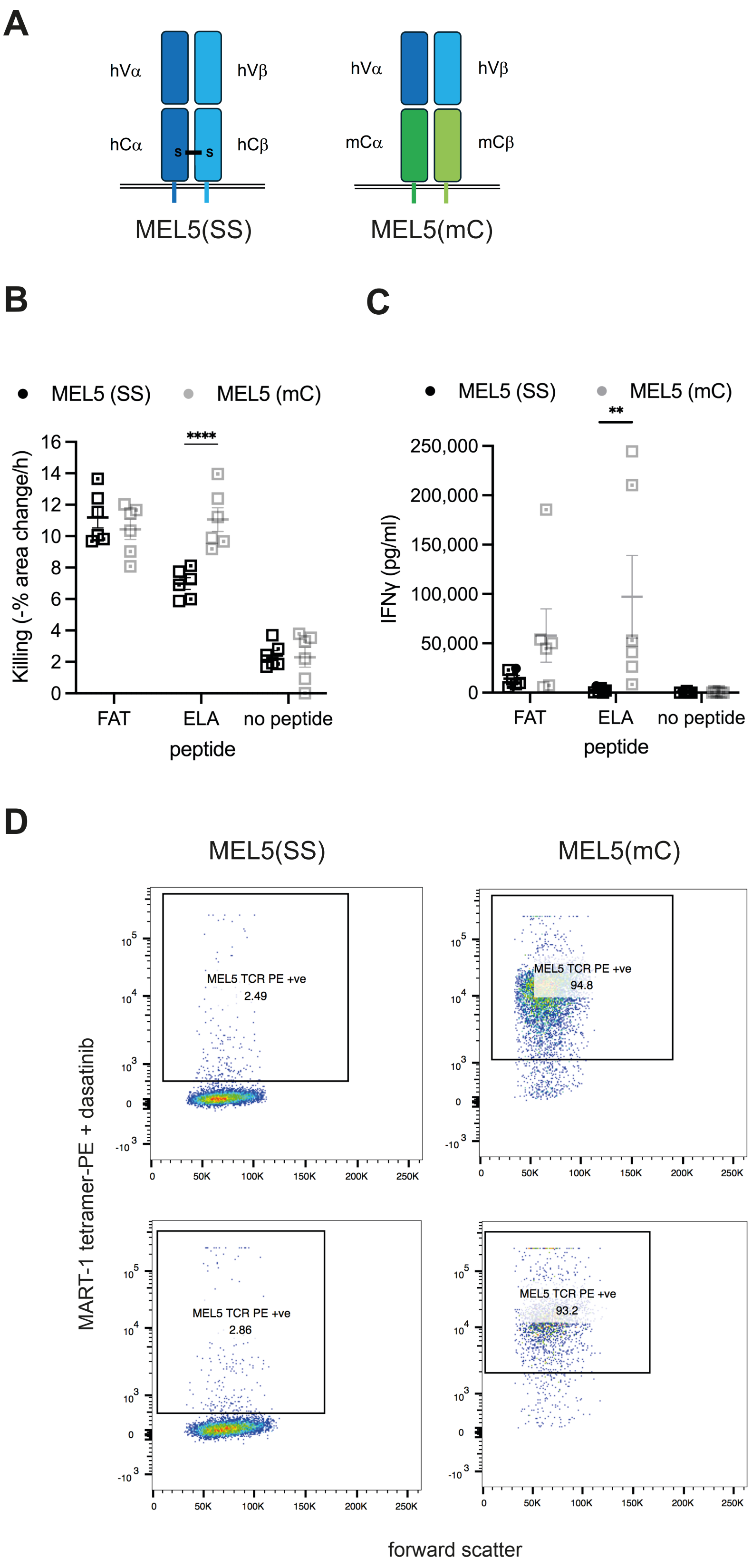


**Stabilization of the MEL5 TCR through murine constant domains is more effective than through an additional disulfide bridge** **A** Schematic of the two versions of the MEL5 TCR. **B, C** Killing of A375 tumor target cells, wild type (open squares) or expressing the folate receptor alpha (open squares with dot, no functional consequences in this experiment) incubated with or without 2µg/ml ELA or FAT MART-1 agonist peptide by CTL transduced to express the MEL5 TCR, stabilized with a disulfide bridge (SS) or murine constant domains (mC) as indicated and IFNγ amounts in the supernatants after 16h interaction as mean ± SEM. 6 independent experiments. Statistical significance determined by paired Two-way ANOVA. B Killing, C IFNγ amounts. **D** Tetramer staining of CTL expressing the indicated version of the MEL5 TCR. 2 independent experiments. ** $p<0.01$, **** $p<0.0001$.

### Figure S5

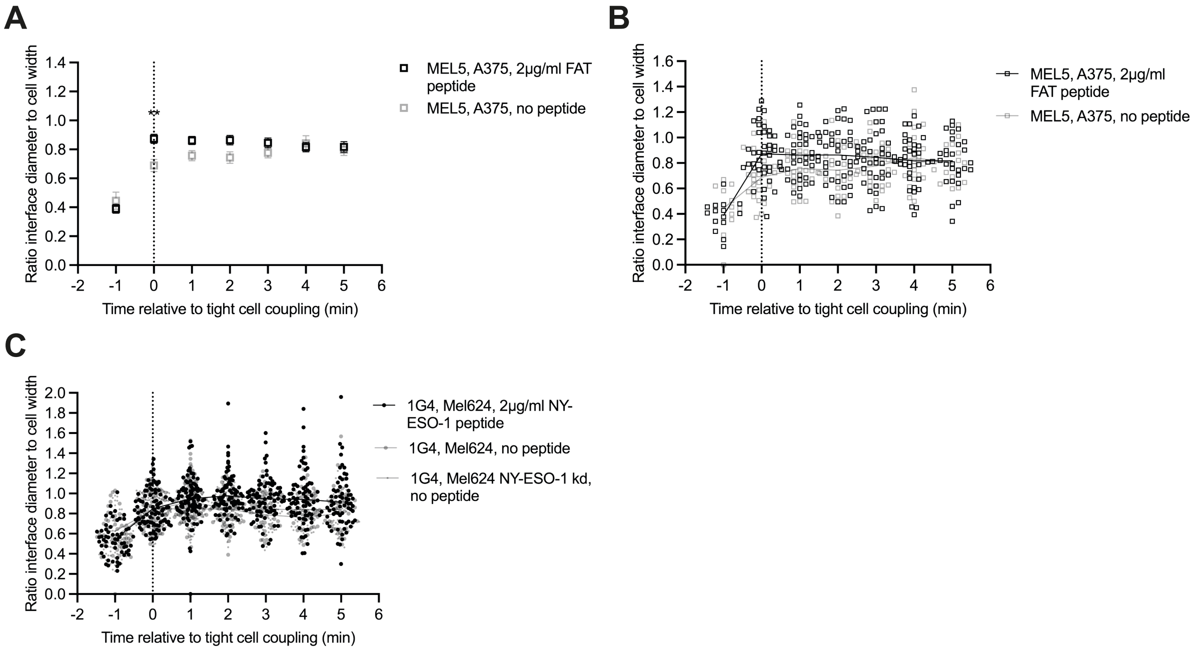


**CTL target cell interface diameters are regulated by stimulus strength in a graded fashion**
**A** Interface diameter relative to the CTL width of MEL5 CTL interacting with A375 cells in the presence of the given amount of the indicated agonist peptide. The MEL5 CTL used in these imaging experiments also express a chimeric costimulatory receptor in the absence of any ligand. 2 independent experiments. Single cell data in B. Statistical significance determined by Two-way ANOVA. **B** Single cell data from B. 40 (FAT peptide) and 21 (no peptide) cell couples analyzed. **C** Single cell data for Fig. 5E. On average 88 (65-101) cell couples analyzed per condition from three independent experiments. ** $p<0.01$

**Figure S6**

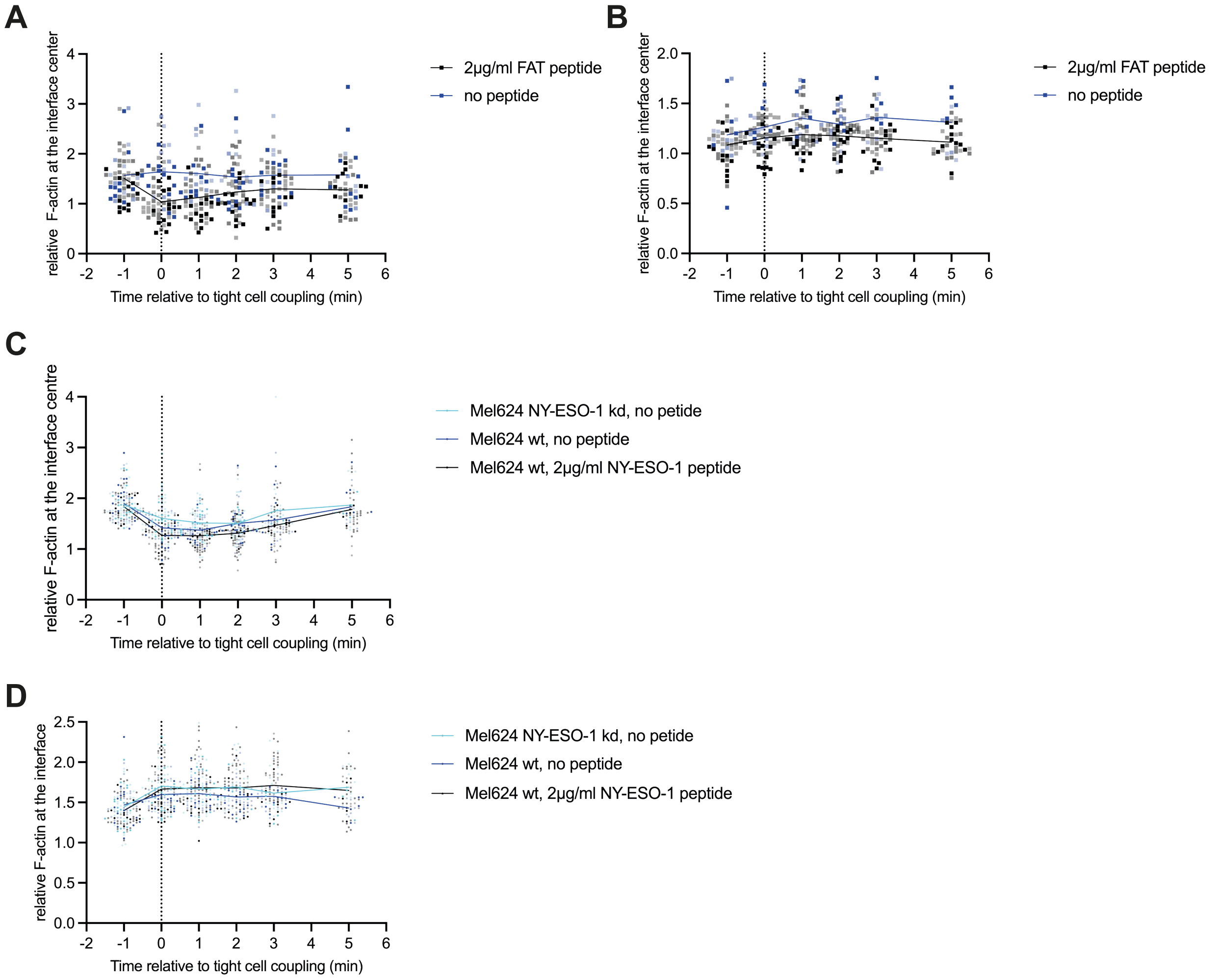


**Only effective cytolysis is associated with F-actin clearance at the center of the CTL target cell interface**

Single cell data for Fig. 6B-E. Independent experiments are indicated by color intensity. On average 45 (21-60) cell couples analyzed per condition. **A** Data for Fig. 6B. **B** Data for Fig. 6C. **C** Data for Fig. 6D. **D** Data for Fig. 6E.

**Figure S7**

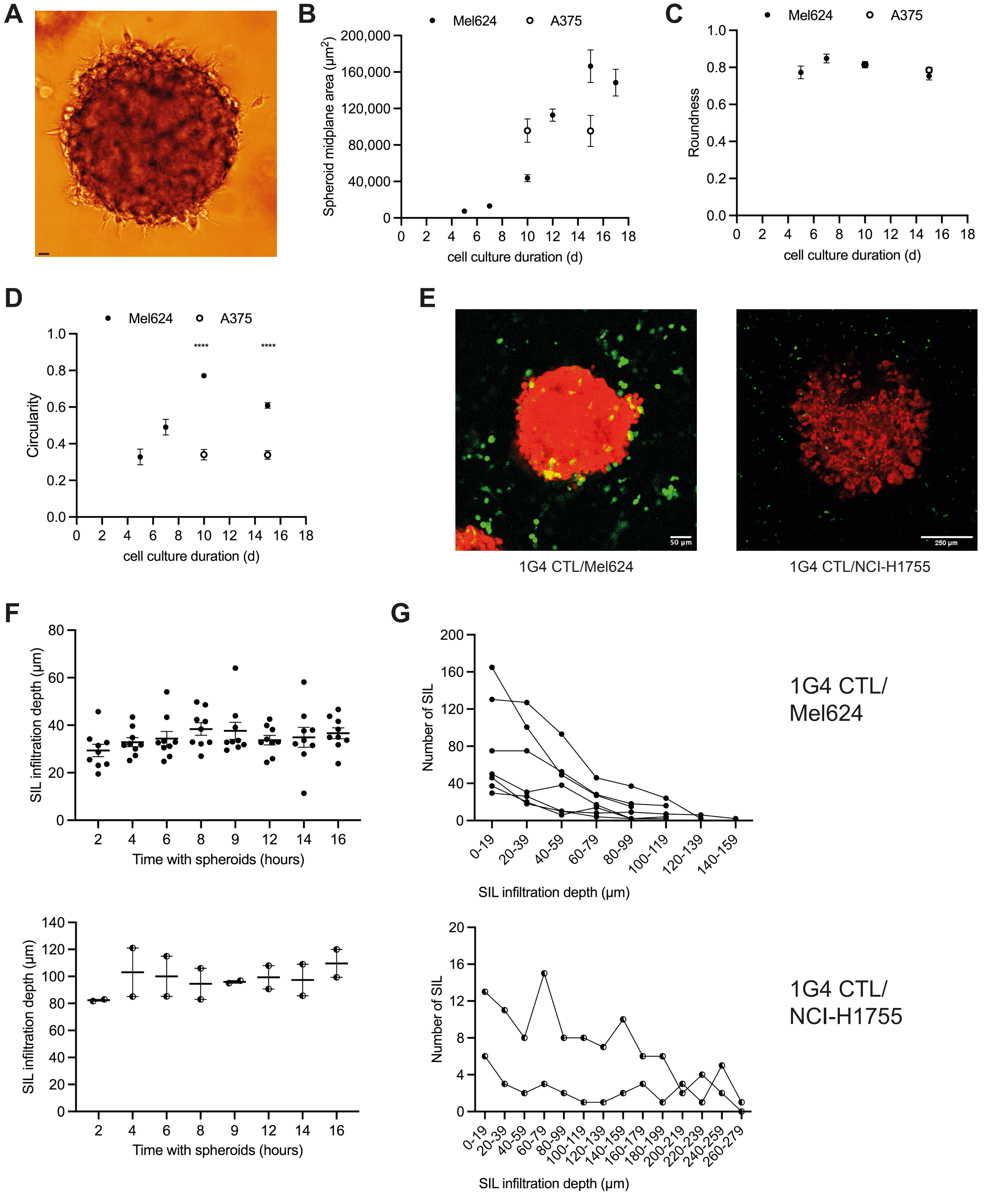


**CTL effectively infiltrate tumor cell spheroids**

**A** Representative image of a Mel624 tumor cell spheroid. Scale bar=20µm. **B-D** Representative growth experiments of Mel625 and A375 spheroids as (B) midplane area, (C) roundness and (D) circularity as mean ± SEM. 4-19 spheroids analyzed on days 5, 7, 12 and 17. 50 spheroids analyzed on days 10 and 15. Statistical difference in circularity at days 10 and 15 determined by Student's t-tests. **E-G** Quantification of 1G4 CTL infiltration into Mel624 or NCI-H1755 spheroids. E Representative images. 1G4 CTL express GFP, tumor cells tdTomato. F Average infiltration depth over time of 1G4 T cells interacting with spheroids as mean ± SEM. G Number of 1G4 T cells at the given infiltration depth window at the 16h time point. 9/2 independent experiments with a total of 36/6 spheroids analyzed. **** $p<0.0001$.

**Figure S8**

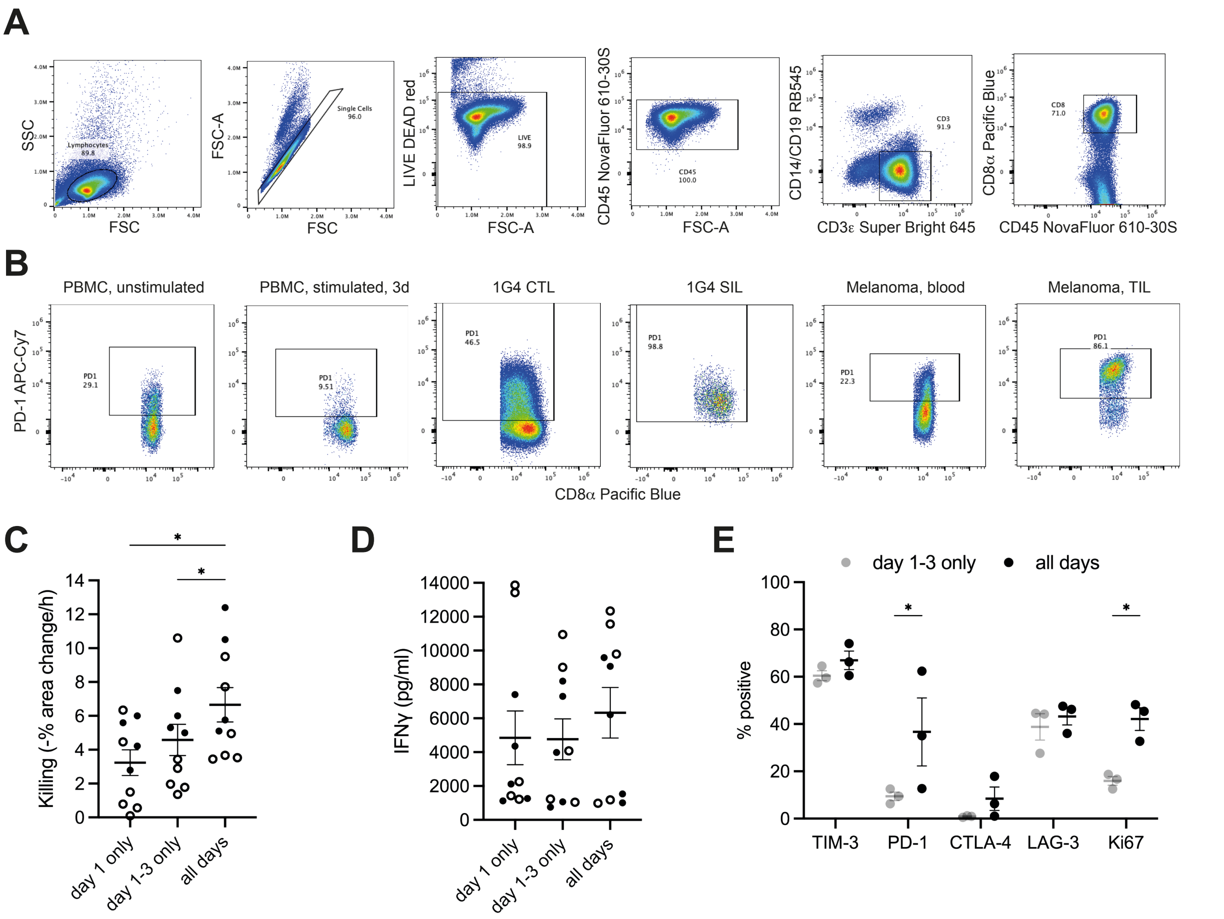


**CTL interaction with spheroids induces an exhausted CTL phenotype**

**A** Sequential gating strategy for the identification of $CD8^+$ T cells. **B** Representative anti-PD-1 staining data are given for the six experimental conditions. Only events gated as CD8α-positive are shown.1 of 3-9 independent experiments. **C, D** Killing of A375 (open symbols) or Mel624 (closed symbols) tumor target cells incubated with 2μg/ml NY-ESO-1 agonist peptide by 1G4 CTL generated in 7-day tissue culture with exposure to anti-CD3/CD28 beads for day 1 only, day 1-3 only or the entire culture period as indicated and IFNγ amounts in supernatants after 16h interaction as mean ± SEM. 10 independent experiments. Statistical significance determined by One-way ANOVA. C Killing, D IFNγ amounts. **E** Percentage of 1G4 CTL, generated in 7-day tissue culture with exposure to anti-CD3/CD28 beads for day 1-3 only or the entire culture period as indicated, positive for the indicated markers as determined by flow cytometry as mean ± SEM. 3 independent experiments. Statistical significance determined by Two-way ANOVA. * $p<0.05$.

### Figure S9

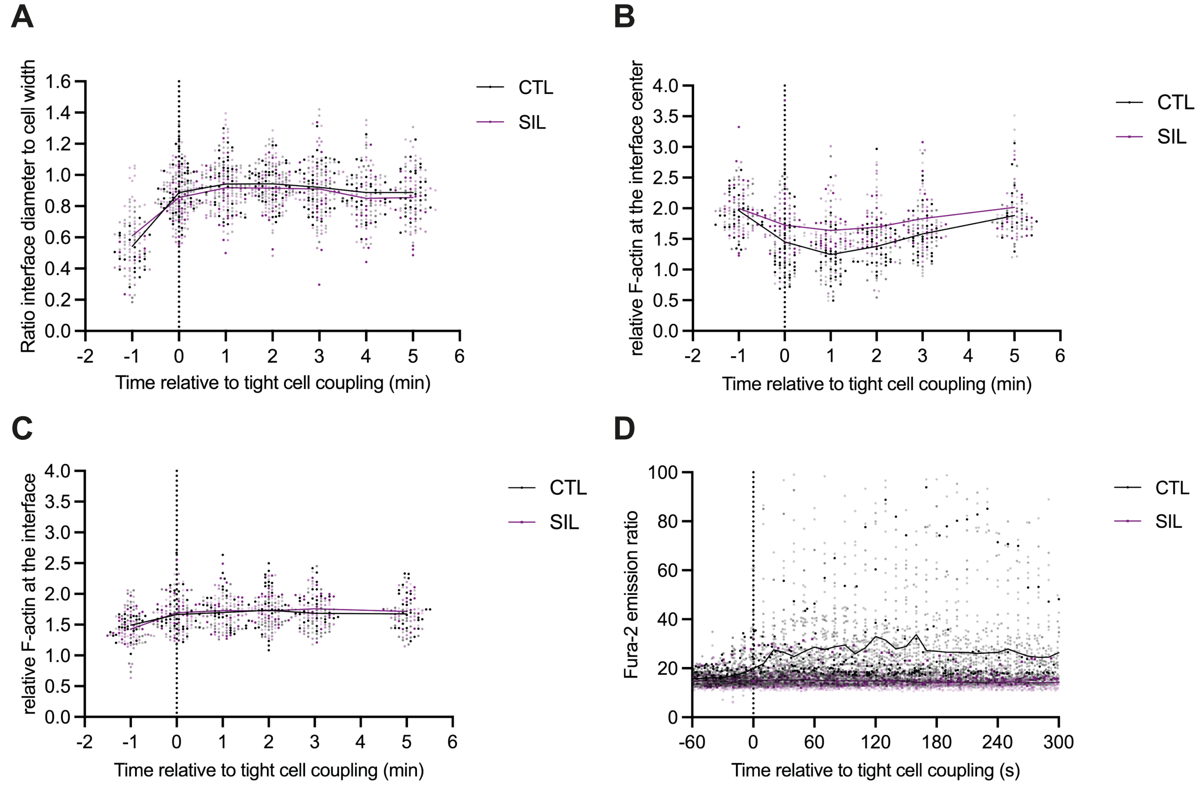


**SIL tumor target cell couples are ill maintained**

**A** Single cell data for Fig. 9F. Independent experiments are indicated by color intensity. 112 (CTL) and 91 (SIL) cell couples analyzed. **B, C** Single cell data for Fig. 9G, H, respectively. Independent experiments are indicated by color intensity. 77 (CTL) and 62 (SIL) cell couples analyzed. **D** Single cell data for Fig. 9I. Independent experiments are indicated by color intensity. 121 (CTL) and 41 (SIL) cell couples analyzed.
